# Stimulus prior and reward probability differentially affect response bias in perceptual decision making

**DOI:** 10.64898/2026.02.16.706079

**Authors:** Christina Koß, Jan Hendrik Blanke, Luis de la Cuesta-Ferrer, Frank Jäkel, Maik C. Stüttgen

## Abstract

Signal detection theory posits that subjects in two-stimulus, two-choice discrimination tasks decide by comparing random samples of an evidence variable to a static decision criterion. While the core assumptions of the theory have received ample experimental support, it has become evident that the decision criterion is not static but subject to trial-by-trial fluctuations and can be influenced by experimental manipulations. The mechanisms governing the trial-by-trial criterion changes are however not well understood. Here, we report results from five experiments in which we subjected rats to a two-stimulus, two-choice auditory discrimination task. In the first three experiments, we investigated the effects of stimulus presentation ratios and reward ratios and provide clear evidence that the effects of changing reward ratios are more pronounced than those of stimulus presentation ratios. A model-based analysis revealed that this effect was due to more than tenfold higher learning rates when reward ratios were manipulated. In two separate experiments, we investigated the effect of reward density (i.e., global reward rate) on criterion learning but failed to find consistent effects. A systematic comparison of three different trial-by-trial criterion learning models based on detection theory, the matching law, and reinforcement learning showed that no model was able to capture the differential effects of stimulus presentation and reward ratios. We conclude that subjects explicitly represent either prior stimulus probabilities or entire stimulus distributions, and accordingly future models need to represent these factors as well.

## 1 Introduction

Signal detection theory (SDT; Green and Swets, 1988) is arguably the standard model for analyzing perceptual decisions. Since its inception in the 1950s (Peterson and Birdsall, 1953; Tanner and Swets, 1954), its influence on psychology and related fields has been steadily increasing and is considered by some as psychology’s “most towering achievement” (Estes, 2002). Today, SDT is being routinely used in fields such as medicine, engineering, neuroscience, and, of course, psychology (for an overview, see Hautus et al., 2022).

One central assumption of SDT is that performance of a subject (observer) in a two-stimulus, two-choice perceptual discrimination task is jointly determined by two factors. The first factor is the degree to which the two stimuli evoke distinct percepts; in SDT, the percept results from sampling from either of two given random variables on a decision axis (a.k.a., “evidence variable”; one distribution for either of the two stimuli). The random variables are usually assumed to be normal and the distance between the means of the two Gaussian distributions, divided by their common standard deviation, is called “sensitivity” or *d^′^*. Second, the observer is then assumed to choose a cutoff criterion along the decision axis, separating evidence samples into two mutually exclusive classes. The decision is taken by comparing the perceived evidence (i.e., the sample from the presented stimulus’ distribution) to an internal decision criterion. Evidence values above the criterion *c* lead to a category 2 response (henceforth, R2), evidence values below to a category 1 response (R1). A central tenet of SDT is the independence of sensitivity and criterion: the former unambiguously reflects the extent to which an observer can distinguish between the two stimuli, the latter however reflects the bias of the observer to emit either of the two responses more or less often. SDT’s core assumptions have been confirmed in a large number of studies (for overview, see Hautus et al., 2022).

SDT tacitly assumes not only static stimulus distributions, but also a static criterion value. However, it was observed early on that this assumption is not tenable (Green, 1964), and there have been numerous demonstrations of fluctuating criterion values since then (e.g., Kubovy and Healy, 1977; Treisman and Williams, 1984; Benjamin et al., 2009; Lak et al., 2020a; Vandevelde et al., 2023). Moreover, ample research has shown that criterion setting is influenced by experimental manipulations, e.g., by unequal stimulus presentation probabilities (SPPs), unequal reward probabilities (RPs), or unequal reward magnitudes for the two responses (Maddox, 2002; Stüttgen et al., 2011, 2013). Nonetheless, SDT does not specify how an observer chooses a certain criterion value in a particular situation. Several different accounts of criterion setting have been proposed over the last decades, but none of them has received widespread support, or even been systematically investigated and compared (for a review, see Hautus et al., 2022). What’s more, most of the criterion setting accounts primarily focus on the steady state, i.e., they do not capture the adaptation of the criterion at a trial-by-trial level but instead merely specify where the criterion will be long after subjects have adapted to a particular experimental condition (e.g., Lee, 1963; Davison and Tustin, 1978; Davison and Jenkins, 1985; Ashby and Maddox, 1990; Busemeyer and Myung, 1992; Davison and Baum, 2000; Maddox, 2002; Brown and Steyvers, 2005; Mueller and Weidemann, 2008; Benjamin et al., 2009; Turner et al., 2011; Killeen et al., 2018).

Models which also specify trial-by-trial updating mechanisms have received less attention (e.g., Luce, 1963; Schoeffler, 1965; Boneau and Cole, 1967; Treisman and Williams, 1984; Erev, 1998; Stüttgen et al., 2011). Building on the seminal work of Kac (1962) and Dorfman and collagues (1971; 1975), we have recently proposed and tested several trial-by-trial accounts of criterion setting (Stüttgen et al., 2011, 2013, 2024; Koß et al., 2024; de la Cuesta-Ferrer et al., 2025). In a series of experiments involving rats performing a single-interval forced choice auditory discrimination task with two or more stimuli and two response options, we found that 1) criterion setting is indeed systematically influenced by reward probabilities, 2) animals adapt their criterion on rewarded but not on unrewarded trials, 3) easily classified stimuli have a smaller effect on criterion updating than difficult stimuli, and 4) steady-state criterion values are determined not by absolute but by relative differences in reward rates (Stüttgen et al., 2024; Koß et al., 2024; de la Cuesta-Ferrer et al., 2025). Moreover, we found indirect evidence that stimulus presentation probabilities influence criterion setting (de la Cuesta-Ferrer et al., 2025). However, none of our previous studies was designed to explicitly characterize the separate and interacting influences of stimulus presentation probabilities, SPPs, and reward probabilities, RPs.

In the present study, we therefore conducted several experiments to investigate the effects of these two factors. Our main research questions are: 1) Do RP and SPP manipulations affect criterion learning to the same extent and in the same way? 2) Does reward density (a.k.a. global reward rate, i.e., the average RP for the two choices), affect the speed of criterion learning, alone or in combination with RP manipulations?

We addressed these questions both through conventional analyses, focusing on steady-state behavior in different experimental conditions, as well as through model-based analyses. For the latter approach, we fit three different trial-by-trial criterion learning models to the data which have performed successfully in past studies, and asked how the models’ fitted parameter values are affected by different experimental manipulations, and which models can fit the choice data and, even more importantly, generate behavior similar to the subjects in these experiments.

The first of the three models is based directly on Kac (1962), Dorfman and Biderman (1971) and Dorfman et al. (1975). Their model assumes that there is a static sensory distribution for each stimulus, and a subject’s decision is taken by comparing the perceived evidence to a criterion. This decision criterion is not static but instead shifts after each trial depending on the feedback: after rewarded responses, the criterion shifts by an amount Δ in the direction that makes the given response more likely subsequently; after unrewarded responses, it shifts in the direction that makes the given response less likely subsequently. Dorfman and Biderman (1971) suggested different model versions, e.g., one where the criterion shifts only after correct (or rewarded) trials, one where it shifts only after incorrect (or unrewarded) trials, and versions which would presuppose that update steps in one or the other direction have the same or different magnitudes. However, neither of their models can reproduce the steady-state behavior that is often observed in animals for conditional discrimination tasks with variable ratio schedules, which has been characterized by the general matching law and its extensions (Davison and McCarthy, 1988, chapter 11). Moreover, de la Cuesta-Ferrer et al. (2025) showed that animals learn from rewards rather than reward omissions, but a model with fixed criterion update steps only after rewarded trials is unstable and will exhibit exclusive choice behavior (see Koß et al. (2024) for a more in-depth discussion). Accordingly, we here use an expanded version of their model which has previously proved its worth (Stüttgen et al., 2013, 2024; de la Cuesta-Ferrer et al., 2025). Briefly, the model posits that the criterion shifts by a fixed step size Δ after every rewarded response into the direction that makes this response more likely to occur in the next trial (see Figure 1AB for illustration). In order to prevent the criterion from drifting off towards positive or negative infinity - a problem identified already by Norman (1974) - we added a leaky integration parameter, *γ*, restricted to values between 0 and 1, by which the criterion is discounted on every rewarded trial. Typically, *γ* is in the range of 0.9-1.0. In the following, we will refer to this model as the “modified KDB model” to distinguish it from previous versions.^1^

**Figure 1:**
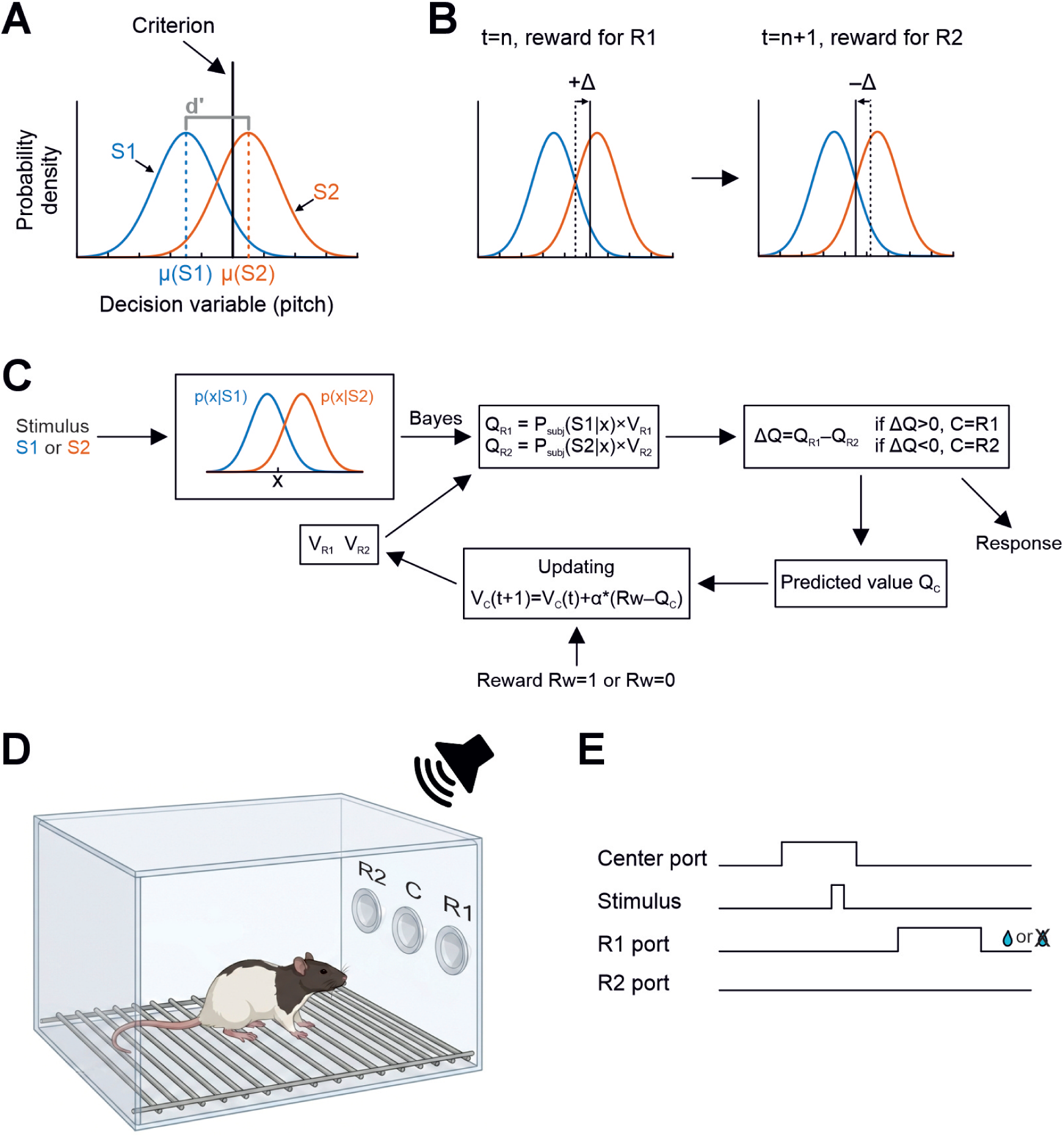
Outline of SDT-based criterion learning models and the behavioral task. **A)** Outline of SDT sensitivity and bias indices. See text for details. **B)** Outline of criterion updating in the KDB and DT models. Left: following rewarded response 1 (R1) in an S1 trial, the criterion shifts to the right with step size Δ, making R1 more likely to occur in the following trial. Right: on the following trial, a rewarded response 2 (R2) in an S2 trial is followed by a criterion shift with step size Δ to the left. Note that Δ is always the same in the KDB model but varies according to the position of the current criterion in the DT model. See Methods for details. **C)** Outline of the RL model. See Methods for details. **D)** Sketch of the behavioral chamber equipped with three conical nose ports. Sound stimuli are played from an overhead speaker. Figure created with FigureLabs. **E)** Diagram of trial epochs. Trials were initiated by the animal continuously poking into the center port for 250-350 ms, followed by presentation of either S1 or S2. Animals responded by either poking into the right (R1) or the left port (R2). Correct responses (S1-R1, S2-R2) were intermittently rewarded by water delivery at the same port, incorrect and unrewarded responses were followed by time out punishment.

The second model is based on the work of Davison and Tustin (1978) who combined SDT and Herrnstein’s (1961) matching law into a description of steady-state choice behavior. Their model, which we refer to as the Davison-Tustin (DT) law, states that the log ratios of the two responses for each stimulus depend linearly on the log ratio of the rewards received for each response:

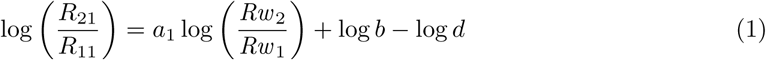

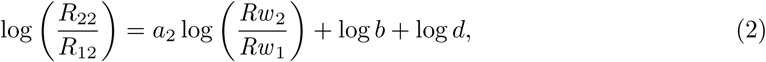

where *R_ji_*denotes the proportion of *R_j_*responses in trials where stimulus *S_i_*was presented and *Rw _j_* denotes the overall proportion of trials where response *R_j_* was emitted and rewarded. It thus features a discrimination index, log *d*, and a bias index, log *b*, both of which are assumed to be independent. Additionally, the parameter *a* specifies the extent to which reward ratios influence the response bias (“sensitivity to reward”). The DT law has received support in several experiments (e.g., McCarthy and Davison, 1979, 1980, 1984). We have shown that the DT law implies behavior according to SDT with logistic stimulus distributions, instead of Gaussian distributions, and a criterion that is proportional to the log ratio of rewards the animal receives for the two responses (Koß et al., 2024). However, the DT law does not provide a trial-by-trial updating mechanism but is fitted only to steady-state choice data, averaged over several experimental sessions which are obtained when performance shows only little fluctuations. We have therefore developed such a model (see Koß et al., 2024). Similar to the KDB model, the DT model assumes that there is a static sensory distribution for each stimulus, and a subject’s decision is taken by comparing the perceived evidence to a criterion that shifts after each rewarded trial by an amount Δ in the direction that makes the given response more likely subsequently. Importantly, in the DT model the size of the update step Δ depends on the response emitted in that trial and on the current criterion position. This dependence is chosen to guarantee that there is a stable equilibrium, and the criterion position in that equilibrium is in line with the DT law.

The third and last of the three models features is based on a combination of SDT and reinforcement learning and is highly similar to a model proposed by Lak et al. (2020b). Unlike the KDB and DT models, it does not directly assume a dynamic criterion, but instead posits that animals store an “action value” V for each possible response. These action values are multiplied with the sensory confidence that the stimulus on the present trial belongs to the respective response category; their product is called Q-value. The animal then emits the response with the higher Q-value, and the action value of the chosen response is updated according to a reward prediction error multiplied with the learning rate *α*. The free parameters in this model are therefore the stimulus variance (monotonically related to *d^′^* in the other models) and the learning rate *α*. The model is sketched in Figure 1C.

To address our research questions, we studied the effects of asymmetric SPPs and RPs both in isolation (Experiments 1 & 2) as well as in combination (Experiment 3). Additionally, we investigated the effect of reward density (Experiment 4) and its interaction with asymmetric reward probabilities (Experiment 5), because reward density has previously been found to affect to the speed of behavioral adaptation (Wittmann et al., 2020).

## 2 Methods

### 2.1 Animals

Nine male rats (Charles River), six weeks old at the time of arrival at the facility, served as experiment subjects. Rats were housed in groups of three in standard type-4 cages in a ventilated cabinet, allowing for continuous monitoring and control of humidity and temperature. Animals were kept on an inverted light-dark cycle (lights off at 8 a.m.). A few days after arrival at the facility, they were handled for at least three days to become habituated to the experimenter. Animals were water-restricted throughout the entire experiment, with water being available in the setup as reward for correct responses. On weekends (Friday through Sunday), water was provided at libitum. If animals did not receive sufficient water during testing, additional water was supplied after the experimental session. All animals continuously gained weight over the course of the experiments (*>*6 months). After conclusion of the experiments, the animals were re-homed to private owners. All subjects were kept and treated in accordance with German guidelines for the care and use of animals in science and conducted in agreement with directive 2010/63/EU of the European Parliament. All experimental procedures conducted with rats were approved by the local ethics review board of the state of Rhineland-Palatinate, Germany (Landesuntersuchungsamt Rheinland-Pfalz, Az. G19-1–094).

### 2.2 Apparatus and procedure

Testing was conducted in standard operant testing chambers (Med Associates) situated in sound-attenuating cubicles. The side wall featured three conical nose ports as response devices (see Figure 1D). Each nose port was equipped with a light beam which was interrupted when the rat poked its nose into the port. Additionally, the two lateral ports were connected to water pumps and featured small wells through which water rewards could be delivered (30 µl per reward). The trial outline is depicted in Figure 1E. Each trial began with the rat poking continuously into the center port for a minimum duration of 0.25 to 0.35 s (randomly sampled from a uniform distribution). Following this initiation interval, one of two auditory stimuli (S1 or S2, see below) was presented. Withdrawal during the initiation interval or during stimulus presentation led to an aborted trial. Aborted trials were not repeated and were not included in the analysis. After stimulus presentation, the rat had 4 s to move to either of the two lateral choice ports to indicate its perceptual decision. Poking into the right choice port (R1) was considered correct after presentation of S1, poking into the left choice port (R2) was considered correct after presentation of S2. Correct responses were rewarded according to the experimental schedule in effect (see below). Incorrect as well as correct but unrewarded responses were followed by a 4-s time out during which house lights went off and no new trial could be initiated. Auditory stimuli consisted of chords composed of 12 pure tones (all having the same amplitude) spaced logarithmically around a center frequency (CF), spanning the frequency range from CV/1.2 to CF*1.2 (Jaramillo and Zador, 2014). All chords were 100 ms in duration and calibrated to a sound pressure level of 74 dB RMS. Rats were trained on the task with two chords spaced far apart (S1: CF=5118 Hz, S2: CF=14774 Hz). After animals mastered the basic task, we gradually decreased the spacing of the two chords until performance was stable with *d^′^* between 1.5 and 2. These two chords were chosen individually for each rat and remained the same until the five experiments were finished. All sounds were generated in Matlab, output from a power 1401-3 AD unit (Cambridge Electronic Design) at a sampling rate of 200 kHz, fed through standard hi-fi amplifiers and played from a loudspeaker mounted on the ceiling of the sound-attenuating cubicle.

### 2.3 Experimental schedules

We conducted five experiments with nine rats. Subjects 1 through 5 served in Experiments 1 through 3, subjects 6 through 9 in Experiments 4 and 5. In Experiment 1, we confronted animals with asymmetric SPPs *π*_1_ and *π*_2_ (taking values of 0.2 vs. 0.8, 0.33 vs. 0.67, 0.5 vs. 0.5, 0.67 vs. 0.33, and 0.8 vs. 0.2), while RPs *ρ*_1_ and *ρ*_2_ were held constant at 0.5 throughout (Table 1A). In the following, we will refer to these conditions by their respective stimulus presentation ratios (SPRs or *π*_1_:*π*_2_), which were 1:4, 1:2, 1:1, 2:1, and 4:1, respectively. In Experiment 2, we held SPPs constant at 0.5 but introduced asymmetric RPs, yielding reward ratios (RRs) which mirror the SPRs in Experiment 1 (Table 1B). In Experiment 3, we directly pitted SPR and RR manipulations against each other, such that in each condition SPR and RR were of the same but opposing magnitudes; e.g., when SPR was 1:4, RR was 4:1. Importantly, if SPR and RR manipulations had the same effects on behavior, their effects should cancel out (Table 1C). In Experiment 4, we kept the RR at 1:1 but varied RPs such that animals experienced different degrees of reward density (or global reward rate). In four conditions, RPs assumed values of 0.25, 0.5, 0.75, and 1 (Table 1D). Finally, in Experiment 5, we varied both RR and reward density to investigate their interaction (Table 1E). Each experimental condition was maintained for at least six consecutive sessions. In all experiments, the two conditions with an inverted ratio, e.g., 1:2 and 2:1, always follow each other directly, and the order of and within these condition pairs was counterbalanced between subjects. For technical reasons, rat 7 was not subjected to condition 5 in Experiment 5.

**Table 1:**
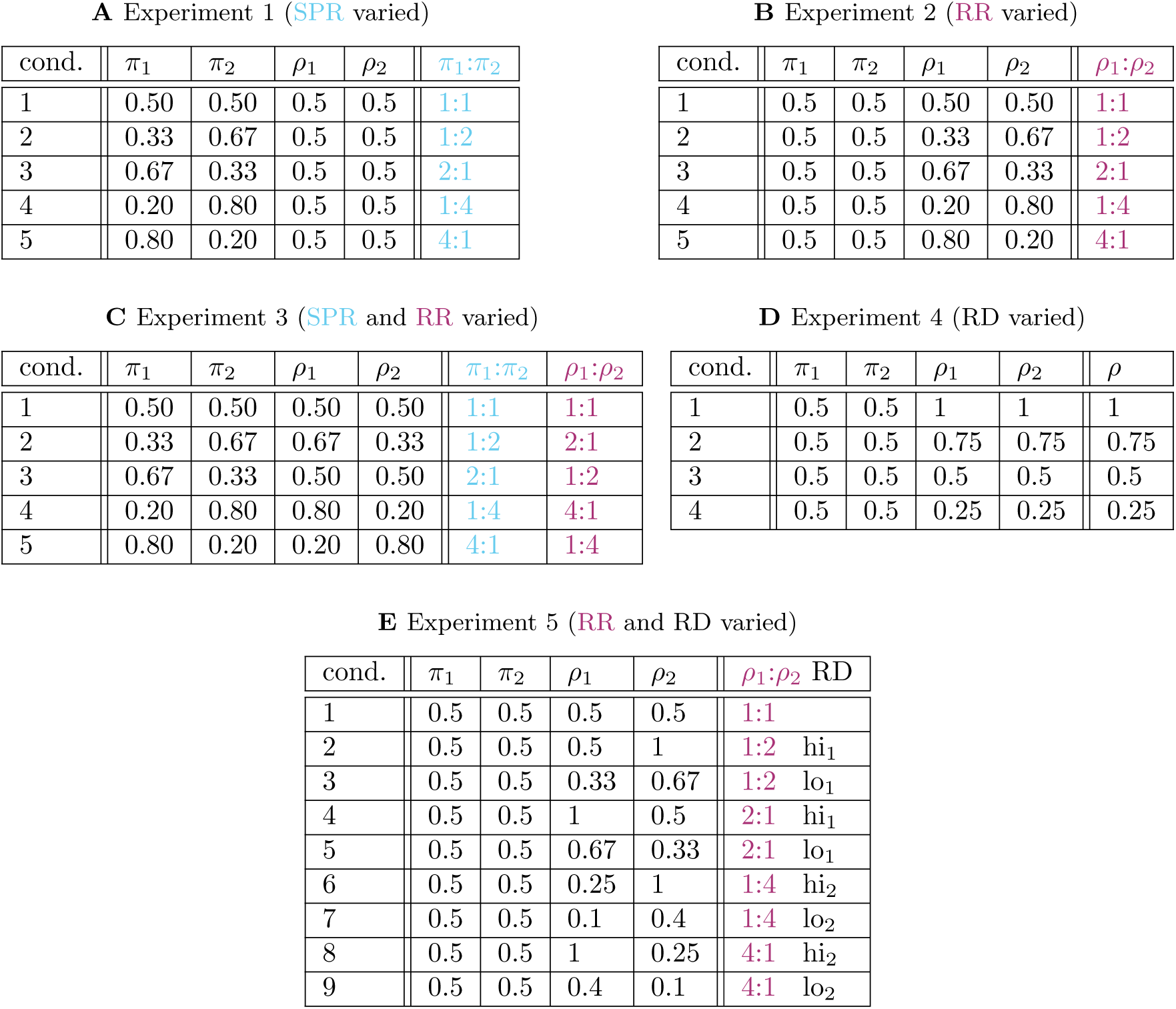
Overview of experimental setups. For each condition, stimulus probabilities *π*_1_, *π*_2_ as well as reward probabilities *ρ*_1_ and *ρ*_2_ are given. The last column(s) summarize each experiment’s relevant manipulation(s).

| cond. | $\pi_1$ | $\pi_2$ | $\rho_1$ | $\rho_2$ | $\pi_1:\pi_2$ |
| --- | --- | --- | --- | --- | --- |
| 1 | 0.50 | 0.50 | 0.5 | 0.5 | 1:1 |
| 2 | 0.33 | 0.67 | 0.5 | 0.5 | 1:2 |
| 3 | 0.67 | 0.33 | 0.5 | 0.5 | 2:1 |
| 4 | 0.20 | 0.80 | 0.5 | 0.5 | 1:4 |
| 5 | 0.80 | 0.20 | 0.5 | 0.5 | 4:1 |

**B** Experiment 2 (RR varied)
| cond. | $\pi_1$ | $\pi_2$ | $\rho_1$ | $\rho_2$ | $\rho_1:\rho_2$ |
| --- | --- | --- | --- | --- | --- |
| 1 | 0.5 | 0.5 | 0.50 | 0.50 | 1:1 |
| 2 | 0.5 | 0.5 | 0.33 | 0.67 | 1:2 |
| 3 | 0.5 | 0.5 | 0.67 | 0.33 | 2:1 |
| 4 | 0.5 | 0.5 | 0.20 | 0.80 | 1:4 |
| 5 | 0.5 | 0.5 | 0.80 | 0.20 | 4:1 |

**C** Experiment 3 (SPR and RR varied)
| cond. | $\pi_1$ | $\pi_2$ | $\rho_1$ | $\rho_2$ | $\pi_1:\pi_2$ | $\rho_1:\rho_2$ |
| --- | --- | --- | --- | --- | --- | --- |
| 1 | 0.50 | 0.50 | 0.50 | 0.50 | 1:1 | 1:1 |
| 2 | 0.33 | 0.67 | 0.67 | 0.33 | 1:2 | 2:1 |
| 3 | 0.67 | 0.33 | 0.50 | 0.50 | 2:1 | 1:2 |
| 4 | 0.20 | 0.80 | 0.80 | 0.20 | 1:4 | 4:1 |
| 5 | 0.80 | 0.20 | 0.20 | 0.80 | 4:1 | 1:4 |

**D** Experiment 4 (RD varied)
| cond. | $\pi_1$ | $\pi_2$ | $\rho_1$ | $\rho_2$ | $\rho$ |
| --- | --- | --- | --- | --- | --- |
| 1 | 0.5 | 0.5 | 1 | 1 | 1 |
| 2 | 0.5 | 0.5 | 0.75 | 0.75 | 0.75 |
| 3 | 0.5 | 0.5 | 0.5 | 0.5 | 0.5 |
| 4 | 0.5 | 0.5 | 0.25 | 0.25 | 0.25 |

Table 1: **Overview of experimental setups.** For each condition, stimulus probabilities $\pi_1$ , $\pi_2$ as well as reward probabilities $\rho_1$ and $\rho_2$ are given. The last column(s) summarize each experiment’s relevant manipulation(s).
| cond. | $\pi_1$ | $\pi_2$ | $\rho_1$ | $\rho_2$ | $\rho_1:\rho_2$ | RD |
| --- | --- | --- | --- | --- | --- | --- |
| 1 | 0.5 | 0.5 | 0.5 | 0.5 | 1:1 |  |
| 2 | 0.5 | 0.5 | 0.5 | 1 | 1:2 | hi<sub>1</sub> |
| 3 | 0.5 | 0.5 | 0.33 | 0.67 | 1:2 | lo<sub>1</sub> |
| 4 | 0.5 | 0.5 | 1 | 0.5 | 2:1 | hi<sub>1</sub> |
| 5 | 0.5 | 0.5 | 0.67 | 0.33 | 2:1 | lo<sub>1</sub> |
| 6 | 0.5 | 0.5 | 0.25 | 1 | 1:4 | hi<sub>2</sub> |
| 7 | 0.5 | 0.5 | 0.1 | 0.4 | 1:4 | lo<sub>2</sub> |
| 8 | 0.5 | 0.5 | 1 | 0.25 | 4:1 | hi<sub>2</sub> |
| 9 | 0.5 | 0.5 | 0.4 | 0.1 | 4:1 | lo<sub>2</sub> |

### 2.4 Analysis

All analyses were run in Python 3.11. The data-sets and the analysis code are available at the OSF project site https://osf.io/fc7pk/.

#### 2.4.1 One-criterion-per-session (OCPS) model

To visualize the animals’ behavior, we show the proportion of R2 in each session. When SPPs are equal for both stimuli, this also represents the animals’ response bias: a bias to emit R2 leads to *P* (*R*2) *>* 0.5 and a bias to emit R1 leads to *P* (*R*2) *<* 0.5. When SPPs are asymmetric, however, P(R2) is confounded with the SPPs because a higher/lower proportion of S2 trials inevitably leads to a higher/lower proportion of R2 responses even when the response bias does not change. The SDT criterion, on the other hand, is unaffected by SPP: a criterion *c <* 0 resp. *c >* 0 always indicates a bias to emit R2 resp. R1, regardless of SPPs. The criterion cannot be directly observed though, so in order to determine the criterion for each session, we fit a standard equal-variance Gaussian SDT model with a fixed discriminability throughout the experiment and one criterion per session as follows.

Let us denote the criterion in session j with *c_j_*. The probability of choosing R2 in session j in a trial where stimulus i was presented is

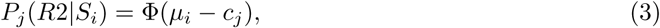

where Φ denotes the cumulative normal distribution. With a fixed criterion per session, these probabilities stay constant throughout the session. Therefore, the expected proportion of R2 responses in the *S_i_* trials in session j is *R*_2*i,j*_ = *P_j_*(*R*2|*S_i_*), and we can replace *P_j_* with the respective proportion. This is equivalent to

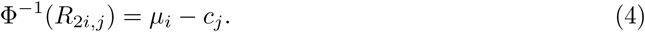

Plugging in the means of the stimulus distributions, *µ*_1_ = −*d^′^/*2 and *µ*_2_ = *d^′^/*2, we get

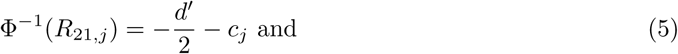

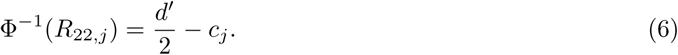

For an experiment with n sessions, this is a model with n+1 parameters: *d^′^* and *c*_1_ to *c_n_*. We determined the stimulus-wise response proportions *R*_21,*j*_ and *R*_22,*j*_ for each session, yielding 2*n data points in total, and fitted the model to these data with a linear least squares regression.

#### 2.4.2 DT law fits

As outlined in Introduction, behavioral data in two-stimulus, two-choice conditional discrimination tasks is commonly found to adhere to the DT law. To validate this assumption for the data from our experiments, we fitted the DT law to the data of each experiment. We restrict the DT law to have the same sensitivity to reward *a*_1_ = *a*_2_ =: *a* for both stimuli and no inherent bias towards either of the responses, i.e., log *b* = 0, for consistency with the DT model, which makes the same assumptions (see Section 2.5.2). Since the DT law only holds in the steady state, we use only the data from the last two sessions of each condition. For each condition, we determined the proportions of R2 and R1 for each stimulus (*R_ji_* denotes the proportion of *R_j_* in trials with *S_i_*), as well as the overall proportions of trials with a rewarded R1 (*Rw* _1_) and with a rewarded R2 (*Rw* _2_), in the last two sessions of the condition. We then fitted the following two straight lines to these data with the method of least squares:

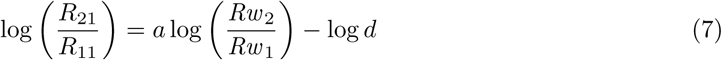

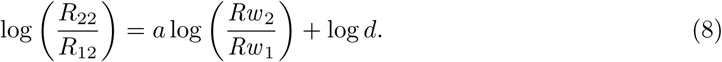

### 2.5 Trial-by-trial modeling

We compared three different models in their abilities to a) fit experimental choice data, and b) generate choices which are qualitatively similar to those of the experimental subjects in forward simulations. Explicitly testing this generative ability mentioned in b) is essential for assessing a model’s validity, because even a model that fits the data well can fail in this regard (see Corrado et al., 2005, for an illustration). Briefly explained, the goodness of fit is based on the log-likelihood of the data summed over all trials. Since the likelihood in each trial takes into account the previous stimuli, responses and rewards that actually occurred in the data, no matter whether the model also produces a similar response sequence or not, simulations can deviate from the predictions of the model fit. Furthermore, we c) investigated if and how the experimental manipulations affect the models’ fitted parameter values to gain insight into the differential effects of SPR, RR and reward density.

We have investigated two of the three models in our previous work (modified KDB and DT, see below); both are based on signal detection theory. The third model is an adaptation of a model proposed by Lak et al. (2020b) and uses a reinforcement learning framework.

Like the analyses, all model fits and simulations were run in Python 3.11 and the code is available at the OSF project site https://osf.io/fc7pk/.

#### 2.5.1 Modified KDB model

The letters K, D, and B stand for the three authors who proposed an early version of the model (see Kac, 1962; Dorfman and Biderman, 1971; Dorfman et al., 1975). We extended their model to include a leak term *γ* which pulls the criterion back towards a baseline position; this can be thought of as a leaky integration mechanism and is necessary to prevent the criterion from drifting towards positive or negative infinity (Stüttgen et al., 2013). Briefly, the model assumes Gaussian stimulus distributions with means *µ*_1_ = −*d^′^/*2 and *µ*_2_ = *d^′^/*2, and a criterion that gets updated from trial to trial. Subjects start out with a criterion *c*(0) = *c_b_*, representing their inherent response bias, and update it after each reward according to the following formulas:^2^

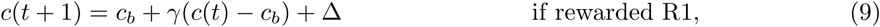

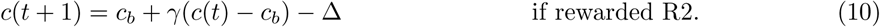

After unrewarded trials, the criterion position remains unchanged at

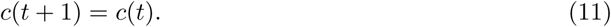

The equilibrium position of the criterion is derived in Appendix A. It depends on the experimental condition, and is influenced by SPP and RP equally. Moreover, it is jointly influenced by the two model parameters Δ and *γ*. The higher the ratio Δ*/*(1 − *γ*), the more extreme the equilibrium criterion will be, for any given experimental condition (see Figure 2A).

**Figure 2:**
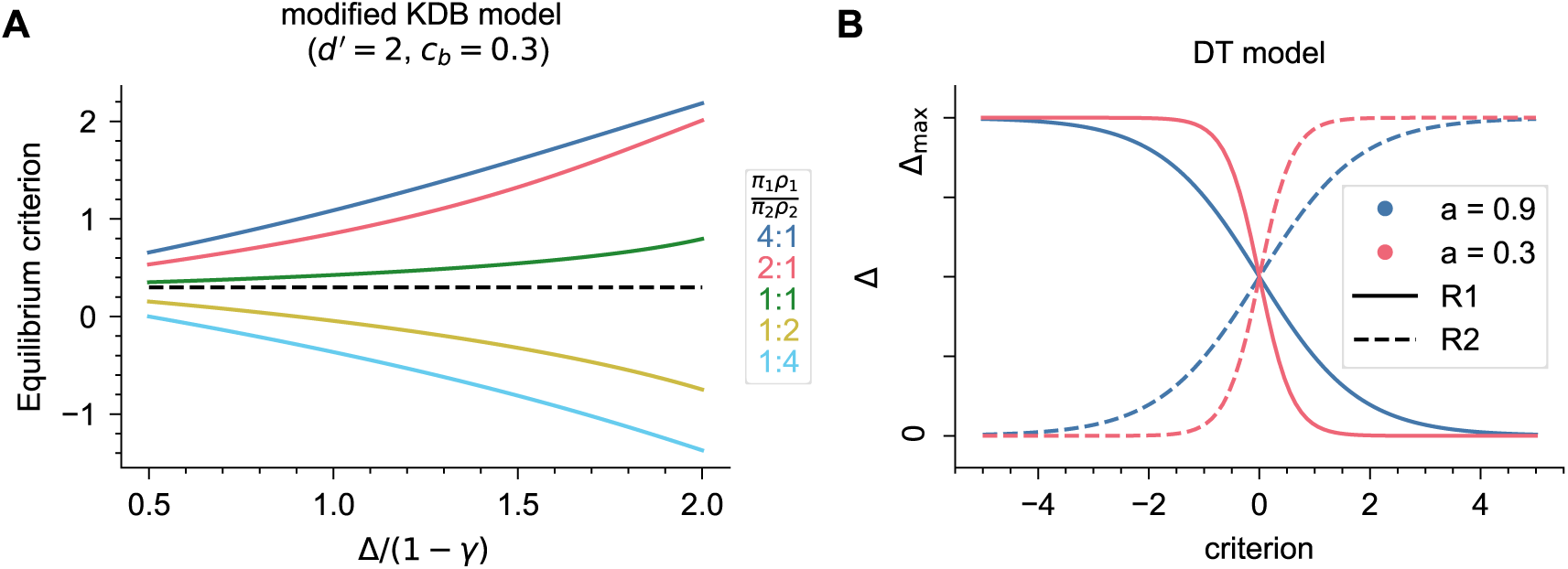
Visualization of model properties. **A)** Equilibrium criterion of the modified KDB model depending on the ratio Δ*/*(1− *γ*) for different experimental conditions, using example values *d^′^* = 2 and *c_b_* = 0.3. The dashed black line is at *c* = *c_b_*, which would be the equilibrium criterion if no rewards were given. **B)** Update step size Δ for the DT model. Δ is shown as a function of the current criterion position *c*, for two different values of the model parameter *a*. Solid lines represent the update step size after a rewarded R1 trial, dashed lines after a rewarded R2 trial.

The model was fitted as described in Stüttgen et al. (2013), by determining the parameters that lead to the maximum likelihood of the data. For a fixed leak term *γ*, the log-likelihood can be expressed as a generalized linear model of the other model parameters *d^′^*, Δ and *c_b_*. Since the log-likelihood is a concave function of these parameters (as Dorfman, 1973, showed), there is a unique maximum, which can be found using standard numerical optimization methods. The model was fitted by repeating this procedure for a range of values of *γ* ∈ (0, 1) and choosing the parameters leading to the overall maximum likelihood.

#### 2.5.2 DT model

The letters D and T stand for the two authors who proposed the original steady-state version of the model (Davison and Tustin, 1978). That model brought together SDT and a very important quantitative relation between choice allocation and experienced reward rates, the Matching Law (Herrnstein, 1961). Since their model (the “DT law”) only described steady-state behavior, we modified and extended it to explain how the criterion is updated at a trial-by-trial level, with an equilibrium criterion at which the DT law is fulfilled (Koß et al., 2024). The DT model assumes logistic stimulus distributions with means 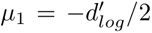 and 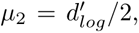 and again a criterion that is updated from trial to trial. In contrast to the modified KDB model, there is no leak term and the size of the update step Δ depends on the current position of the criterion and is different for each response option. Subjects initially use a criterion *c*(0) = *c*_0_, and after each reward they update it according to

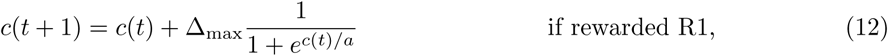

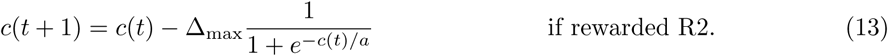

After unrewarded trials, the criterion position remains unchanged at

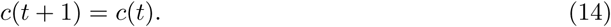

The parameter Δ_max_ is the maximum step size and functions as a constant scaling parameter, and *a* determines the subject’s sensitivity to reward, as in the DT law.

Figure 2B shows how the update step size Δ depends on the criterion for different values of *a*. At *c* = 0, the update steps in either direction have a size of Δ = Δ_max_*/*2. As soon as the criterion gets shifted towards either side of zero, the update steps further in the same direction get scaled up more and more (to a maximum of Δ_max_) whereas update steps back in the opposite direction get scaled down more and more (to a minimum of 0). The lower *a*, the steeper the logistic functions in Equations 12 and 13, which means that when the criterion gets shifted away from *c* = 0, update steps further outwards will be scaled down, and steps back inwards will be scaled up. Therefore, for lower *a* the criterion does not move outwards as fast and as much, resulting in a behavior that is less extreme and less sensitive to the experimental manipulations.

The initial criterion at the start of the experiment, *c*_0_, cannot yield any insights about the experimental manipulations and only has a small influence on the model fits overall, especially for long experiments.^3^ We still include it as a free parameter in the model for the sake of completeness (Dorfman et al., 1975, have shown for a similar model that including *c*_0_ as a free parameter is essential to get valid results).

The model, like the modified KDB model, was fitted by determining the parameters that lead to the maximum likelihood of the data. For each combination of *a*, Δ_max_ and *c*_0_, the log-likelihood can again be expressed as a generalized linear model of 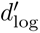 with a unique maximum that can be found using numerical optimization methods. The model was fitted by running a grid search over *a*, Δ_max_ and *c*_0_, and for each combination determining the best 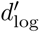 and corresponding maximum log-likelihood. Then, the parameter combination that led to the overall maximum likelihood was chosen. The grid ranges used were

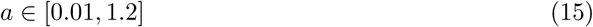

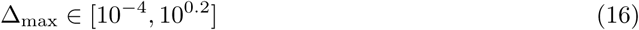

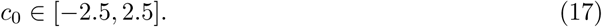

For some experiments, these ranges were extended as some of the found maxima were located at the edge of the smaller ranges: For Experiment 3, the range for *a* went up to 3.0 and for Experiments 4 and 5 up to 2.0. Moreover, for Experiment 4 and 5, the range for Δ_max_ was (10*^−^*^5^, 1).

#### 2.5.3 RL model

Lak and colleagues (2020a; 2020b) have proposed and tested a reinforcement learning model for value-based perceptual decisions, which we adapted and fitted to our data for comparison purposes. The model, just like the modified KDB and DT models, is based on SDT and assumes that perceived evidence comes from a certain distribution dependent on the shown stimulus. The subject keeps track of internal estimates *V*_1_ and *V*_2_ (“action values”), which indicate how much reward it expects from emitting a correct R1 or R2. In each trial, the subject chooses the response for which it expects the higher reward given the perceived evidence. The expected reward for each response *R_i_* is determined as

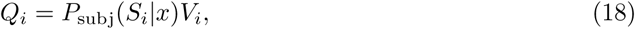

where *P*_subj_(*S_i_*|*x*) denotes the subject’s belief that the perceived evidence *x* comes from stimulus *S_i_* (referred to as “decision confidence”). This belief is computed in a Bayesian manner from the prior belief that stimulus *S_i_* is presented and the likelihood that the perceived evidence occurs in a trial with *S_i_*. This is a computation internal to the subject, therefore, rather than the “real” *P* (*S_i_*) = *π_i_* and *p*(*x*|*S_i_*), the subject’s subjective belief about stimulus probabilities *P*_subj_(*S_i_*) and stimulus distributions *p*_subj_(*x*|*S_i_*) have to be used. It is thus computed as

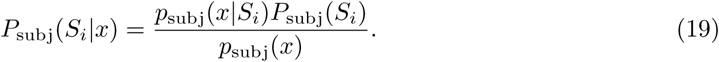

The stimuli employed in our experiments differ greatly from the ones that were used by Lak et al. (2020a,b), so we need to adjust the model assumptions about *p_subj_*(*x*|*S_i_*) and *P_subj_*(*S_i_*) for our setup. Our experiments featured exactly two stimuli, designed to be relatively well distinguishable (around 80 percent correct responses; see Section 2.2). Subjects had extensive experience with these stimuli from being trained on the task before the experiments started. Therefore, our version of the RL model assumes that subjects’ subjective belief about the stimulus distributions matches the real distributions. Still, the subjects cannot know the real SPPs, since these are manipulated in some of the experiments. Our RL model assumes the most uninformative belief of *P_subj_*(*S*1) = *P_subj_*(*S*2) = 0.5, which are also the real SPPs in the experiments without SPP manipulation (Experiment 2, 4 and 5).

With these assumptions,

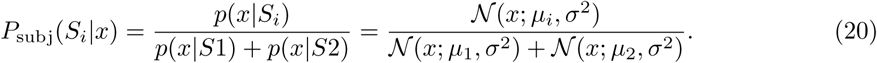

As always in SDT, scaling *σ*^2^ and *µ_i_* by the same factor does not alter the model, so one of the parameters has to be fixed arbitrarily. Often, *σ*^2^ = 1 is chosen, and that is also how we set up the modified KDB model and the DT model. Lak et al. (2020b) instead fixed the stimulus means and kept *σ*^2^ as a free parameter, so we do the same in our RL model by setting *µ*_1_ = −0.5 and *µ*_2_ = 0.5, i.e., fixing the distance between stimulus distributions to 1.

If *Q*_1_ *> Q*_2_, the subject emits R1, and R2 otherwise. After the response has been emitted and either has been rewarded or not rewarded, the subject updates its value estimate for the chosen response, while the estimate for the unchosen response remains unchanged. If response *R_i_*was chosen, *V_i_*is updated according to

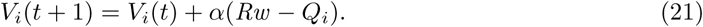

This update is proportional to the difference between actually received and expected reward, i.e., the reward prediction error, with *α* being a learning rate that scales the update size.

For the RL model, computing the likelihood of the data under the model is intractable, so instead we used the same fitting method as Lak et al. (2020b). A grid search was run over *α* and *σ*^2^. For each parameter combination on the grid, 20,000 to 200,000 model simulations were run, using the original sequences of stimuli and potential rewards.^4^ Then, for each trial the proportion of simulated responses that match the response *R_C_* that was chosen by the subject, *P* (*R_C_*), was computed. An alternative “negative log-likelihood” measure NLL*_alt_* was defined as the total 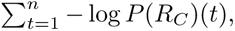 i.e. summed over all *n* trials, to approximate the real 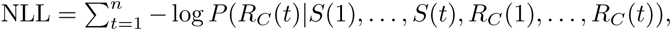 and the best-fitting parameters were determined as the ones with the lowest NLL_alt_. The grid ranges used were

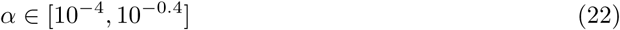

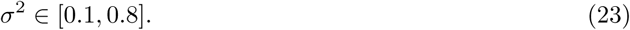

For Experiments 4 and 5, the range for *α* was extended to be *α* ∈ (10*^−^*^5^, 1) as some of the found maxima were located at the edge of the smaller parameter range.

#### 2.5.4 Model versions with multiple learning rates

In order to investigate the effect of reward density on the models’ learning rates, we introduce a version of each model that allows different learning rates (i.e., Δ in the modified KDB model, Δ_max_ in the DT model and *α* in the RL model) for different sets of trials. Trials from sessions that share the same reward density were grouped together. In Experiment 4, this means grouping together trials that belong to the same condition, leading to four trial groups with *r* = 1, 0.75, 0.5 and 0.25 respectively (cp. Table 1D). In Experiment 5, it means grouping together trials in each pair of conditions that share the same (but mirrored) reward probabilities, leading to five trial groups, one for the baseline condition and one for each reward density hi_1_, lo_1_, hi_2_ and lo_2_ (cp. Table 1E). The multiple-learning-rate versions of the models work exactly like the single-learning-rate models described above, just with more learning rate parameters Δ*_g_*, Δ_max*,g*_ or *α_g_*that are used for the criterion update in the trials belonging to trial group *g*. In the following, when we don’t refer to one of the specific models, the learning rate parameter for trial group *g* is denoted by LR*_g_*.

The modified KDB model with multiple learning rates can be fitted with the same procedure as the single-learning-rate version. To fit the DT and RL models, grid searches were employed, again. However, with the number of parameters increasing to up to 8 for the DT model and up to 6 for the RL model (in Experiment 5, where the models have five learning rate parameters), the method described above is computationally not feasible anymore. Fortunately, each learning rate parameter LR*_g_* only affects behavior directly during trial group *g*. Trial groups earlier in the experiment are not affected, and the effect on trial groups later in the experiment is indirect, only via the criterion position at the end of trial group *g* (and thus the beginning of the next trial group). As mentioned before, the influence of the starting criterion on behavior quickly vanishes. Therefore, an approximate sequential fitting procedure can be used, where log-likelihoods are computed only for data from one trial group *g* at a time, with a partial parameter grid including just the one learning rate LR*_g_* together with the non-learning-rate parameters, rather than computing the log-likelihood of the full data set on the full up-to-8-dimensional grid. See Appendix B for details.

### 2.6 Model comparisons

The modified KDB and DT models both use the NLL as a measure of goodness of fit on an individual data-set. Since both models have the same number of parameters, the values can directly be compared for each subject in an experiment to conclude which of the two models provides the better fit to these data. Lower values indicate a better fit.

When comparing the single-learning-rate and multiple-learning-rates versions of each model, the number of parameters varies between versions. Therefore, a direct comparison of NLLs is not informative. Instead, we used the Bayesian information criterion (BIC), which is computed as

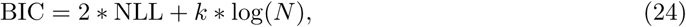

where *k* is the number of free parameters of the fitted model and *N* the number of trials in the data. A lower BIC corresponds to a better fitting model.

For the RL model, the computation of the NLL and thus also the BIC is intractable, and it is unclear how the approximate goodness-of-fit measure used for this model’s fit, NLL_alt_, relates to the “real” NLL. Therefore, the RL models’ NLL_alt_ cannot be compared to the modified KDB and DT models’ NLL. However, comparisons between the single-learning-rate and multiple-learning-rate version of the RL model, using an approximate BIC_alt_ measure that is computed from the NLL_alt_ according to Equation 24, are still appropriate, because that measure relies on the same approximation in both model versions.

## 3 Results

### 3.1 Experiments 1 and 2

Experiment 1 and 2 were designed to study the effects of different stimulus probability ratios and reward ratios, respectively. In Experiment 1, the reward probabilities for both stimuli were held constant at 0.5, while the stimulus probability ratio (SPR) varied (see Table 1A). In Experiment 2, both stimuli were presented with equal probabilities of 0.5 while the reward ratio (RR) varied (see Table 1B). Importantly, SPRs and RRs had the same set of values in both experiments; therefore, if SPRs and RRs exert identical effects, we would expect to observe highly similar behavior in the two experiments. Also, the two manipulations predict identical optimal (i.e., reward maximizing) criteria, because the optimal criterion is located where the likelihood-ratio 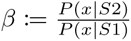 is equal to 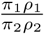 (Green and Swets, 1988, chapter 1.7.2).

The behavior of an example subject is shown in Figure 3A (Experiment 1) and B (Experiment 2). As expected, asymmetric ratios reliably produced a response bias toward the response designated correct for the more frequent stimulus (Experiment 1) or the stimulus associated with a higher reward probability (Experiment 2). In Figure 3AB, this bias is evident in the proportion of responses which shifted towards the preferred response already in the first session after a change in contingency (marked by vertical dashed lines).

**Figure 3:**
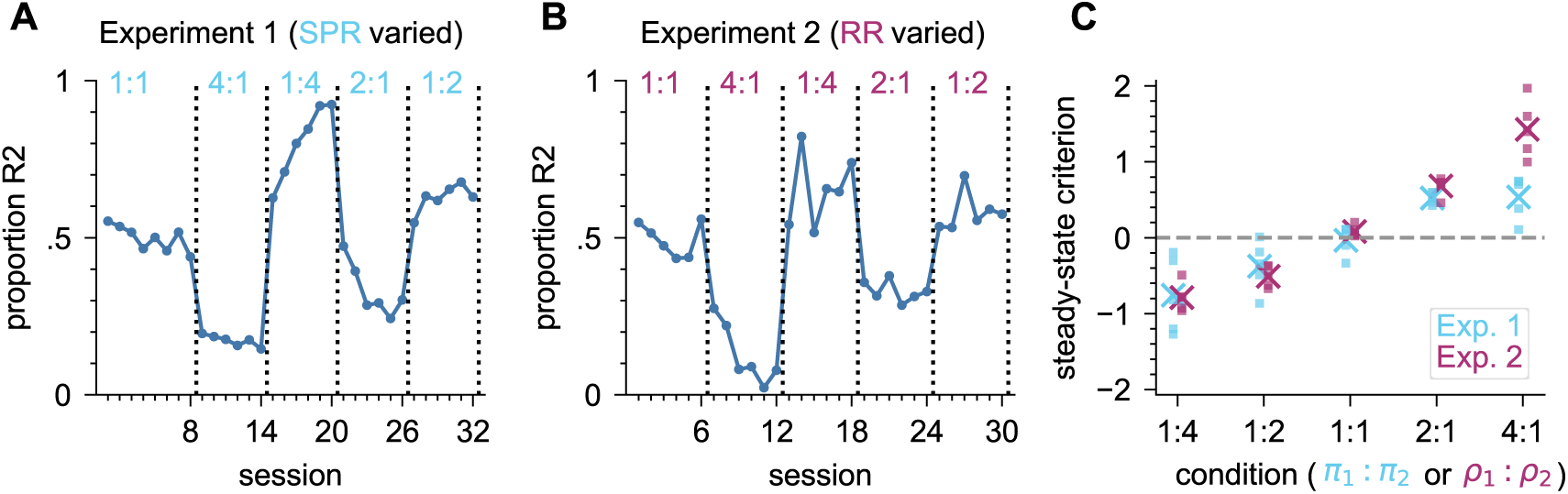
Behavior in Experiment 1 (SPR varied) and 2 (RR varied). **A)** Response proportions in Experiment 1 for example subject 1. Each data point is the proportion of R2 in one session. **B)** As A) but for Experiment 2. **C)** Steady-state criterion for all subjects, i.e., criterion from the one-criterion-per-session model averaged over the last two sessions per condition. Each data point stems from one experimental condition, and means across animals are shown as an ×. Conditions are ordered from SPR (Experiment 1) or RR (Experiment 2) most strongly favoring R2 towards most strongly favoring R1.

To investigate the animals’ steady-state behavior, the proportion of R2 is not informative, as it is confounded with the stimulus presentation probabilities and therefore does not allow for meaningful comparisons between the two experiments. The SDT decision criterion, however, is by definition independent of SPPs. Accordingly, we obtained an SDT decision criterion value for each session by fitting a one-criterion-per-session model to the data, i.e., a standard SDT approach where we assume a fixed *d^′^* throughout the experiment and a fixed criterion in each session (see Section 2.4.1). A bias to emit R1 manifests in positive criterion values, while a bias for R2 manifests in negative criterion values. The steady-state criteria of all animals for both experiments are depicted in Figure 3C. As conditions got more biased towards one of the two response options, so did the animals’ behavior, which can be seen in the absolute value of the criterion becoming more extreme. Note that the reward manipulations in Experiment 2 had a slightly stronger effect on the criterion than the stimulus manipulations in Experiment 1. For the same ratio of *ρ*_1_ : *ρ*_2_ or *π*_1_ : *π*_2_ respectively, the steady-state criteria tended to be more extreme in Experiment 2, where the bias is induced by varying the reward ratio; this effect was small, but visible in every single condition.

#### DT law fits

As outlined in Introduction, we expect that subjects’ behavior adheres to the DT law. Also, the DT model predicts steady-state behavior in line with the law, and the steady-state behavior generated by the modified KDB model with realistic parameter values also follows it closely enough that they are hard to distinguish empirically. We therefore fitted the DT law to the data (for details, see Section 2.4.2), to see if the data in our experiments is in line with this established law and to investigate potential differences between the animals’ behavior in the two setups. Results can be seen in Figure 4. The DT law fits yield two behavioral measures: The sensitivity to reward *a* (slope of the lines) measures how strongly the animals’ response ratio depends on the ratio between received rewards for either side, and the discriminability log *d* (distance between the S1- and S2-lines) is a measure of how well animals can distinguish between the stimuli. While the DT law generally held in both experiments when considered separately, subjects were more sensitive to variations in received reward when reward probabilities were varied than when stimulus probabilities were varied: *a* is consistently higher in Experiment 2 (RR varied) than in Experiment 1 (SPR varied) for every subject. While log *d* is also slightly higher in Experiment 2 (RR varied) than in Experiment 1 (SPR varied), the magnitude of this effect is very small and barely visible in the graphs.

**Figure 4:**
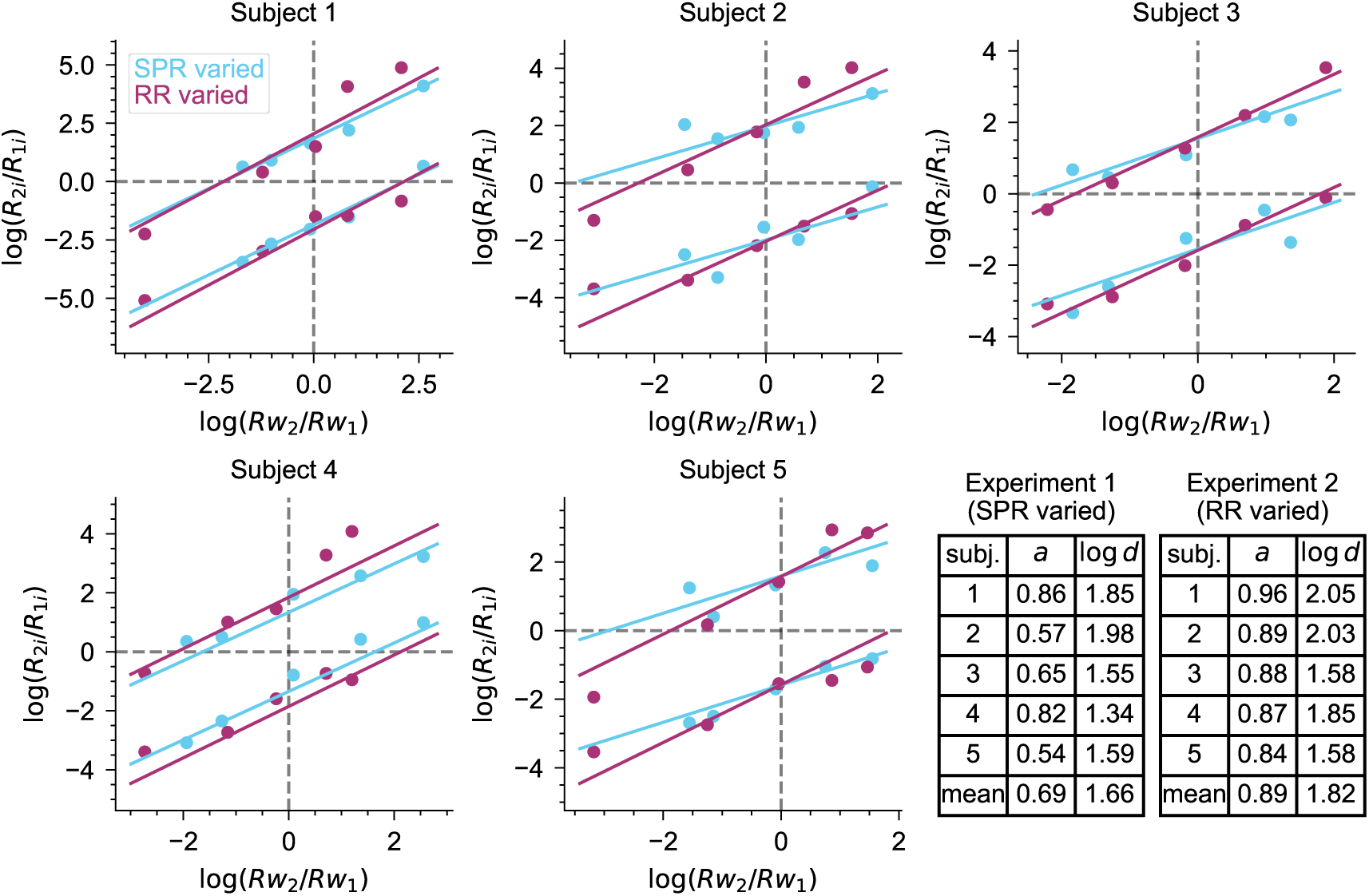
DT law fits for Experiment 1 (SPR varied, blue) and 2 (RR varied, purple). Stimulus-wise log response ratio is plotted against reward ratio for each subject. Each data point stems from one experimental condition. The lower and upper lines show the fitted linear relationship for stimulus 1 and 2 trials respectively. The fitted parameter values are shown in the bottom right corner.

#### Model fits

Next, we investigated if the animals’ trial-by-trial choices can be explained by the three models (modified KDB model, DT model, and RL model), and whether the models also reveal differences between the two experiments. We fitted each model to the data from both experiments (for more details on the fitting procedures, see Sections 2.5.1, 2.5.2 and 2.5.3), and additionally ran 100 model simulations for each model, using the best-fitting parameters and subjecting the model to the same stimulus and potential reward sequences as the original subjects. Figure 5 shows the proportion of R2 for each session for the original data, the model fit and the simulations, for both experiments for an example subject. Plots for all subjects can be found in the appendix in Figures C.2–C.6 (one figure for each experiment and model).

**Figure 5:**
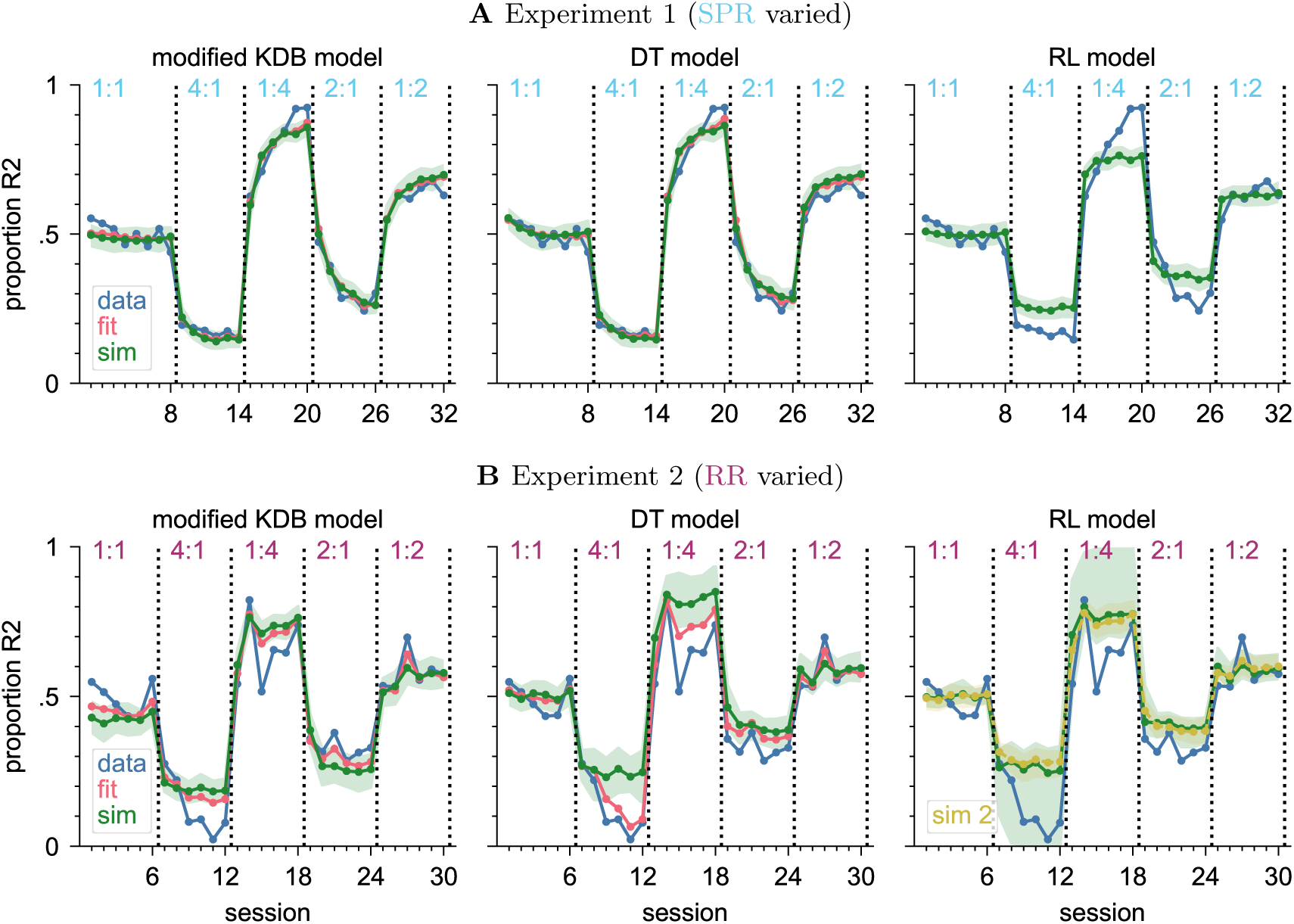
Model fit and simulations applied to the data of example subject 1 in Experiment 1 and 2. **A)** Experiment 1 (SPR varied). **B)** Experiment 2 (RR varied). Each data point is the proportion of R2 in one session. Blue line: data. Red line: proportion of R2 predicted by the model fit (using the best-fitting parameters), i.e., session average of P(R2) in each trial given the actual trial history. Green line: median of 100 simulations (using the best-fitting parameters), shaded green area: central 95% of the simulations. Yellow: same as green, but for the parameters at the NLL minimum, in case these differ from the best-fitting parameters (see footnotes 6, 5 and 7).

The modified KDB and DT models’ behavior generally followed the subjects’ behavior closely, though there are some conditions where the two deviate (Figure 5, left and middle columns). For example, the two models (by design) cannot capture situations where a subject behaves asymmetrically in some conditions. In this example, this is the case for Experiment 2 in the 1:4 and 4:1 conditions, which feature identical, only flipped, reward ratios, but the subject’s behavior was much more shifted towards R1 in the 4:1 condition. Also, the variance in the data exceeds the variability of the simulations: Several data points lie outside of the central 95% of the simulations. This means that there are other sources of behavioral noise that are not reflected in either model, in which the only source of noise is the perceptual noise from the stimulus.

To investigate which model fits the data better, Table 2 compares the NLL for each subject and experiment (see Section 2.6). For Experiment 1, both models fitted equally well, with each model being a better fit for some of the subjects. Experiment 2 (RR varied) was overall better fitted by the DT model, which has a much lower NLL for all subjects except subject 3, where the modified KDB model fits slightly better. Still, both models captured the behavior relatively well in both experiments, especially taking into account that not only the goodness of fit is relevant, but also the behavior in simulations. E.g., for the example subject shown in Figure 5B, in the 4:1 condition the fit of the DT model (middle column) predicts response proportions closer to those in the data than the modified KDB model (left column). However, this is not reflected in the model simulations, which stay equally far away from the data for both models.

**Table 2:** NLL comparison between the modified KDB model and the DT model for Experiment 1 and 2, using the respective best fitting parameters for each subject. The lower NLL (better fit) is highlighted in green for each subject.

| Experiment 1 (SPR varied) |  |  | Experiment 2 (RR varied) |  |  |
| --- | --- | --- | --- | --- | --- |
| subj. | modified KDB | DT | subj. | modified KDB | DT |
| 1 | 5936 | 5938 | 1 | 5649 | 5604 |
| 2 | 5140 | 5135 | 2 | 4803 | 4752 |
| 3 | 5440 | 5429 | 3 | 4854 | 4860 |
| 4 | 5934 | 5965 | 4 | 6037 | 5997 |
| 5 | 6294 | 6290 | 5 | 6732 | 6709 |

For the RL model (Figure 5, right), we cannot show predicted response proportions for the model fit, since these require the calculation of P(R2) in each trial given the history of presented stimuli and issued rewards, which is not tractable (see Section 2.5.3). Therefore, we show only the median and central 95% interval of 100 simulations. Judging by the visual impression, the simulations of the RL model seem to reproduce the subjects’ behavior in Experiment 2 (RR varied) similarly well as those of the other two models: The simulation trajectories generally follow the data, but deviate from them in some conditions, especially where the animals behaved asymmetrically. Also, the variance of the simulations for the RL model is considerably higher than the variance for the other two models, especially in the 1:4 / 4:1 conditions. For Experiment 1 (SPR varied), the behavioral trajectory of the RL model’s simulations is a visibly worse match to the original data than the other models’ simulations. Specifically, the RL model simulations fail to capture the slow, gradual adjustment that follows a condition change, and instead the proportion of R2 very quickly converges to a new steady state. The steady state of the simulations also does not match the actual steady state at the end of the condition. This is presumably related to the previous issue: If the proportion of R2 does not increase slowly but immediately reaches a fixed steady-state level, this level has to be lower than the steady state in the original data in order to also explain the behavior earlier after the condition change reasonably well, where the subject’s behavior has not reached a new steady state yet while the RL model already has.

Comparing the best-fitting parameters of the models – i.e., the ones that yield the lowest negative log-likelihood, which means that the probability of generating the observed data is highest – between the two experiments can give valuable insights into the animals’ behavior.

The best-fitting parameters for the modified KDB model are shown in Table 3.^5^ The fitted *d^′^*, which measures the ability to distinguish between the two stimuli, was slightly higher in Experiment 2 (RR varied) than in Experiment 1 (SPR varied), but again the difference was small in terms of expected performance. The magnitude of criterion updates after rewarded trials was much larger in Experiment 2 (RR varied) than in Experiment 1 (SPR varied) for this model, too, because the fitted update step size Δ is an order of magnitude larger. The leak term *γ*, which determines how fast the past criterion updates are “forgotten” by pulling the criterion back to the baseline *c_b_*, and update step size Δ jointly influence the steady-state criterion. All else being equal, a higher ratio 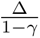 leads to a more extreme steady-state criterion (see Appendix A), which implies a larger behavioral impact of the experimental manipulation. Hence, we calculated this ratio for each subject. In Experiment 2 (RR varied), the ratio 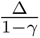 was always larger than in Experiment 1 (SPR varied). This is evidence that animals’ response behavior is more sensitive to reward probability manipulations than to stimulus probability manipulations.

**Table 3:** Best-fitting parameters of the modified KDB model for Experiment 1 (blue) and 2 (purple). In parentheses are the parameters that yield the lowest NLL in the case where these do not explain the data well and therefore the parameters corresponding to a second minimum are used as the best fit (cp. footnote 5).

| subj. | $\gamma$ | | $d'$ | | $\Delta$ | | $c_b$ | | $\frac{\Delta}{(1-\gamma)}$ | |
| --- | --- | --- | --- | --- | --- | --- | --- | --- | --- | --- |
|  | E1 | E2 | E1 | E2 | E1 | E2 | E1 | E2 | E1 | E2 |
| 1 | 0.995 | 0.964 | 2.05 | 2.17 | 0.0048 | 0.052 | 0.04 | 0.21 | 0.60 | 1.44 |
| 2 | 0.998 | 0.951 | 2.22 | 2.38 | 0.0018 | 0.064 | -0.01 | 0.16 | 0.90 | 1.31 |
| 3 | 0.994<br>(0.192) | 0.887 | 1.67<br>(1.76) | 1.93 | 0.0046<br>(0.341) | 0.138 | 0.05<br>(0.09) | 0.03 | 0.77<br>(0.42) | 1.22 |
| 4 | 0.992 | 0.933 | 1.45 | 1.99 | 0.0086 | 0.084 | -0.10 | -0.04 | 1.07 | 1.25 |
| 5 | 0.995 | 0.957 | 1.70 | 1.86 | 0.0036 | 0.055 | 0.05 | 0.19 | 0.72 | 1.28 |
| mean | 0.995 | 0.938 | 1.82 | 2.07 | 0.0047 | 0.079 | 0.00 | 0.11 | 0.81 | 1.30 |

For the DT model, the best-fitting parameters are shown in Table 4.^6^ Similar to the observations in the DT law fits (see Section 3.1), the fitted *a* in the DT model, which influences how sensitive the response behavior is to variations in received rewards, tended to be higher in Experiment 2 (RR varied), though the effect was less clear here. While the fitted 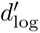 was larger in Experiment 2 (RR varied) for every subject – meaning that subjects can distinguish between the stimuli better there – this effect was behaviorally very small. E.g., for subject 1, the expected baseline performance with optimal criterion setting only increases from 85% correct responses to 87%. This is again in line with the results from the DT law fits (see Section 3.1). The most notable difference between the two experiments is that the decision criterion gets updated by a much larger amount after each rewarded trial in Experiment 2 (RR varied) than in Experiment 1 (SPR varied), since the maximum update step size Δ_max_ was more than 20 times larger there. There is no clear pattern observable that would indicate that the initial criterion position *c*_0_, which reflects the response bias at the beginning of the experiment, differed systematically between the two experiments – unsurprisingly so, since animals cannot know the experimental manipulation before the experiment has started.

**Table 4:** Best-fitting parameters of the DT model for Experiment 1 (blue) and 2 (purple). In parentheses are the parameters that yield the lowest NLL in the case where these do not explain the data well and therefore the parameters corresponding to a second minimum are used as the best fit (cp. footnote 6).

| subj. | $a$ | | $d'_{\log}$ | | $\Delta_{\max}$ | | $c_0$ | |
| --- | --- | --- | --- | --- | --- | --- | --- | --- |
|  | E1 | E2 | E1 | E2 | E1 | E2 | E1 | E2 |
| 1 | 0.81 | 0.89 | 3.49 | 3.79 | 0.0191 | 0.282 | -0.48 | -1.49 |
| 2 | 0.99 | 0.97 | 3.79 | 4.27 | 0.0062 | 0.295 | -0.39 | -0.23 |
| 3 | 0.67<br>(0.05) | 0.83 | 2.81<br>(3.00) | 3.32 | 0.0140<br>(1.297) | 0.426 | 1.08<br>(-0.16) | -0.97 |
| 4 | 0.82 | 0.81 | 2.43 | 3.40 | 0.0294 | 0.285 | -0.03 | 1.09 |
| 5 | 0.63 | 0.87 | 2.86 | 3.25 | 0.0097 | 0.178 | 0.76 | -1.80 |
| mean | 0.79 | 0.87 | 3.08 | 3.61 | 0.0157 | 0.293 | 0.19 | -0.68 |

The best-fitting parameters for the RL model are shown in Table 5.^7^ There were substantial differences between the two experiments in the learning rate *α*, which scales the updates of the value estimates, and thereby the updates of the resulting decision criterion, after each trial. For every subject, *α* was larger in Experiment 2 (RR varied) than in Experiment 1 (SPR varied) by an order of magnitude or more. On the other hand, the perceptual uncertainty parameter *σ*^2^ was smaller in Experiment 2 (RR varied), implying better perceptual discriminability of the stimuli. But these differences, while they may appear substantial, were again rather small in terms of behavioral significance. E.g., for subject 1, the expected baseline performance with optimal criterion setting only increased from 83% correct responses to 85%, which is very similar to the results from the fits of the modified KDB and DT models.

**Table 5:** Best-fitting parameters of the RL model for Experiment 1 (blue) and 2 (purple). In parentheses are the parameters that yield the lowest NLL in the cases where these do not explain the data well and therefore the parameters corresponding to a second minimum are used as the best fit (cp. footnote 7).

| subj. | $\sigma^2$ | | $\alpha$ | |
| --- | --- | --- | --- | --- |
|  | E1 | E2 | E1 | E2 |
| 1 | 0.27 | 0.23<br>(0.33) | 0.010 | 0.20<br>(0.032) |
| 2 | 0.22 | 0.18 | 0.005 | 0.25 |
| 3 | 0.40<br>(0.34) | 0.26 | 0.009<br>(0.18) | 0.18 |
| 4 | 0.68 | 0.28 | 0.009 | 0.11 |
| 5 | 0.37 | 0.30<br>(0.39) | 0.005 | 0.14<br>(0.014) |
| mean | 0.39 | 0.25 | 0.008 | 0.18 |

We fitted the DT law to simulated data for all models to see if the models capture the same regularities between response ratios and reward ratios as the experimental data (see Appendix C.2). For the modified KDB model and the DT model, this is indeed the case, as the simulated data could be reasonably fitted and the resulting fits were similar to those of the experimental data. For the RL model, the same was true for Experiment 2 (RR varied). For Experiment 1 (SPR varied), the simulated data could also be reasonably fitted, but they displayed a substantially lower sensitivity to reward *a* than the experimental data. This strengthens the finding that the RL model cannot accurately reproduce the animals’ behavior in the setup with varying stimulus probabilities.

#### Summary

Overall, our analyses showed that the RR manipulations in Experiment 2 biased the animal’s behavior more strongly than the SPP manipulations in Experiment 1.

The differences in steady-state criteria were small but consistent across all conditions, and sensitivity to reward as observed in the DT law fits was higher in Experiment 2, when RR was varied, for each subject.

The fits of the three models also showed that animals update their behavior from trial to trial differently in the two experiments. They seem to shift their decision criterion after rewarded trials by a much larger amount (reflected in Δ in the modified KDB model, Δ_max_ in the DT model and *α* in the RL model) in a scenario with unequal reward ratios for the two responses, compared to a scenario with unequal stimulus presentation probabilities. Moreover, the animals seem to be able to distinguish slightly better between the stimuli (reflected in *d^′^* in the modified KDB model, 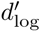 in the DT model and *σ*^2^ in the RL model) when reward probabilities are manipulated, though this effect is so small that it is barely relevant behaviorally. The fits of the modified KDB model and the DT model also strengthen the observation from the DT law fits that animals are more sensitive to changes in received reward in the reward-manipulation setup. In the DT model, this shows up as a larger *a*, and in the modified KDB model as a larger ratio 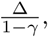 both of which indicate more extreme steady-state behavior.

All three models capture the subjects’ behavior relatively well in Experiment 2 (RR varied). For the modified KDB model and the DT model, the same is true for Experiment 1 (SPR varied), while the RL model does not capture the slow adjustment of response behavior and displays steady-state behavior that is less extreme than the subjects’.

### 3.2 Experiment 3

Experiment 1 and Experiment 2 showed differential effects of stimulus and reward manipulations when either was manipulated in isolation. Experiment 3 combines the two manipulations in an opposing manner. Stimulus probabilities and reward probabilities were pitted against each other such that *π*_1_*ρ*_1_ = *π*_2_*ρ*_2_, i.e., an SPR of 1:2 would occur together with a RR of 2:1, etc. (see Table 1C). Recall that the optimal criterion is located where the likelihood-ratio 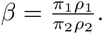 In this experiment, the ratio on the right hand side stays constant at 1, which implies a neutral optimal criterion of *c* = 0.

In Experiments 1 and 2, we found that animals displayed a stronger response bias (higher sensitivity to reward as well as more extreme steady-state criteria) in reaction to varying reward probabilities than to varying stimulus probabilities. If this was also the case when RRs and SPRs are varied simultaneously by the same amount but opposing ratios, as in Experiment 3, one would expect the animals to adapt an overall bias in the direction of the stimulus associated with the higher reward probability, i.e., a decision criterion that is shifted to make the response associated with that stimulus more likely. This bias should be stronger with ratios *ρ*_1_ : *ρ*_2_ and *π*_1_ : *π*_2_ further away from 1.

The behavior of an example subject is shown in Figure 6A. Since this experiment tries to compare the biasing effects of SPR and RR, but the proportion of R2 is not a direct measure of response bias as it is confounded with SPR (see Section 2.4.1), one cannot draw conclusions about the bias from looking at the raw behavior. Therefore, we looked at the criteria resulting from the one-criterion-per-session model. Figure 6B shows the criterion development over sessions for subject 1 (results for the other subjects in Figure D.1 in the appendix). The criterion does indeed deviate markedly from the neutral (and optimal) criterion value of 0 and follows the pattern predicted from the results of Experiments 1 and 2: It takes on negative values (bias towards R2) in the conditions where *ρ*_1_ *< ρ*_2_ and *π*_1_ *> π*_2_, and positive values (bias towards R1) in the conditions where *ρ*_1_ *> ρ*_2_ and *π*_1_ *< π*_2_, and the values tend to be more extreme in the conditions with SPRs and RRs further away from 1. The same pattern holds for the steady-state criteria per condition for all subjects, shown in Figure 6C. This is evidence that reward probability manipulations influence the animals’ response behavior stronger than stimulus probability manipulations, not only in isolation, but also when both are combined.

**Figure 6:**
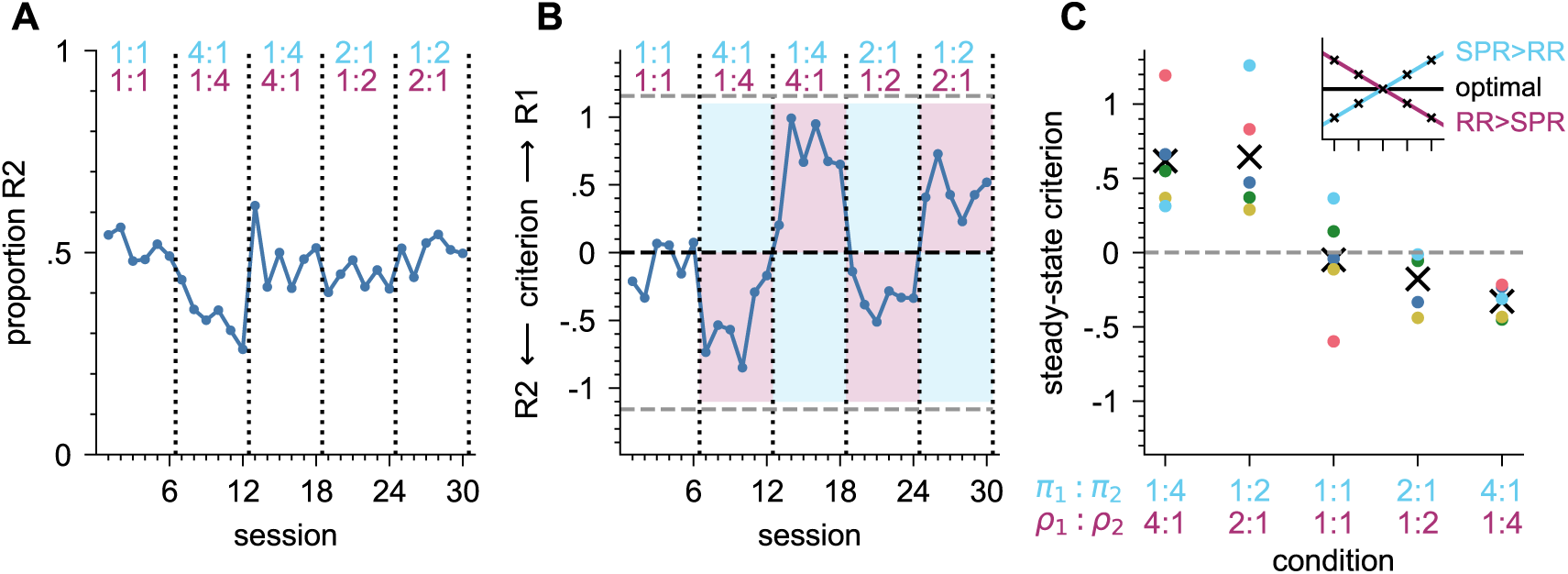
Behavior in Experiment 3 (SPR and RR varied). **A)** Response proportions for example subject 1. Each data point is the proportion of R2 in one session. **B)** Criterion development for example subject 1 as fitted by the one-criterion-per-session model. For reference, zero (black) and the fitted stimulus means ±*d^′^/*2 (gray) are marked as dashed lines. A positive/negative criterion indicates a bias to emit R1/R2 respectively. Blue/purple shading marks the areas where the criterion would indicate that SPR/RR has a stronger effect respectively. **C)** Steady-state criterion for all subjects. Each data point stems from one experimental condition, with colored dots representing individual subjects and means across animals shown as an ×. Conditions are ordered from SPR most strongly favoring R2 and RR favoring R1 towards vice versa. Optimal behavior would result in a slope of 0; a positive/negative slope would indicate that SPR/RR has a stronger effect.

#### Model fits

Again, we fitted the three models to the data and additionally ran 100 simulations for each of them, as we did for the two first experiments. Qualitatively, none of the three models could explain the animals’ behavior in this experiment.

The modified KDB model and the DT model were suitable for fitting the data and generating similar behavior in Experiment 1 and 2, but neither of them could account for the response biases observed in Experiment 3 (see Figure 7, left and middle). The main reason for this inability to reproduce the behavior is that the position of the equilibrium criterion *c_eq_* in both models is determined by 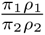 (Koß et al., 2024; de la Cuesta-Ferrer et al., 2025). This ratio has a constant value of 1 throughout Experiment 3. For the DT model, a ratio of 1 corresponds to *c_eq_* = 0 (which is also the optimal criterion here), while the modified KDB model allows a response bias that depends on the model parameters. In both models, the equilibrium criterion stays the same in all conditions. Therefore, they cannot produce the condition-dependent criteria observed in the data, which yield to different response biases (see Appendix D.2 for a more detailed explanation).

**Figure 7:**
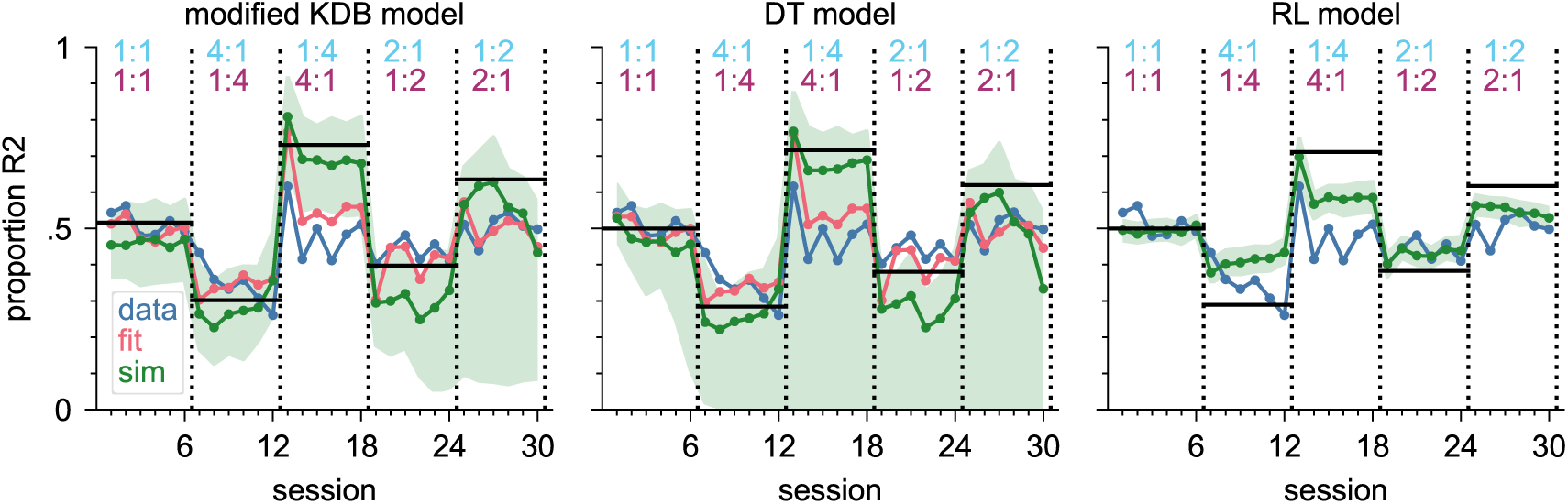
Model fit and simulations applied to the data of example subject 1 in Experiment 3. Each data point is the proportion of R2 in one session. Blue line: data. Red line: proportion of R2 predicted by the model fit (using the best-fitting parameters), i.e., session average of P(R2) in each trial given the actual trial history. Green line: median of 100 simulations (using the best-fitting parameters), shaded green area: central 95% of the simulations. Black lines: proportion of R2 that would result from a fixed criterion at *c_eq_* (modified KDB model) or *c* = 0 (DT and RL models) as a reference.

The RL model provided a reasonable fit to Experiment 2, but not to Experiment 1. It also could not reproduce the animals’ behavior in Experiment 3 (see Figure 7, right, for an example subject and Figure D.2 in the appendix, right column, for the other subjects). This is unsurprising, because Experiment 3, like Experiment 1, includes manipulations of SPR and therefore the experimental conditions deviate from the RL model’s stimulus prior with *π*_1_ = *π*_2_ = 0.5.

In summary, the results from Experiment 3 support our conclusions from Experiments 1 and 2 that 1) RR manipulations bias behavior more strongly than SPR manipulations and 2) therefore, none of the models can satisfactorily generate behavior that resembles that of the animals in a scenario where both RR and SPR are manipulated.

### 3.3 Experiment 4

Experiment 4 investigated the influence of reward density on behavior. Both stimulus probabilities and reward probabilities were kept symmetric at a 1:1 ratio, and only the overall reward density was varied by changing the reward probabilities from condition to condition (see Table 1D). In this experiment, the optimal criterion is again *c* = 0 in every condition. As expected, all animals indeed maintained criterion values close to 0 throughout the experiment, resulting in response proportions close to 0.5 (see example subject in Figure 8A; the other subjects’ data is included in Figure E.1 in the appendix). Figure 8B shows steady-state criterion values for all subjects. While the steady-state criteria deviated somewhat from 0, no systematic relationship between reward probability and steady-state criterion is apparent.

**Figure 8:**
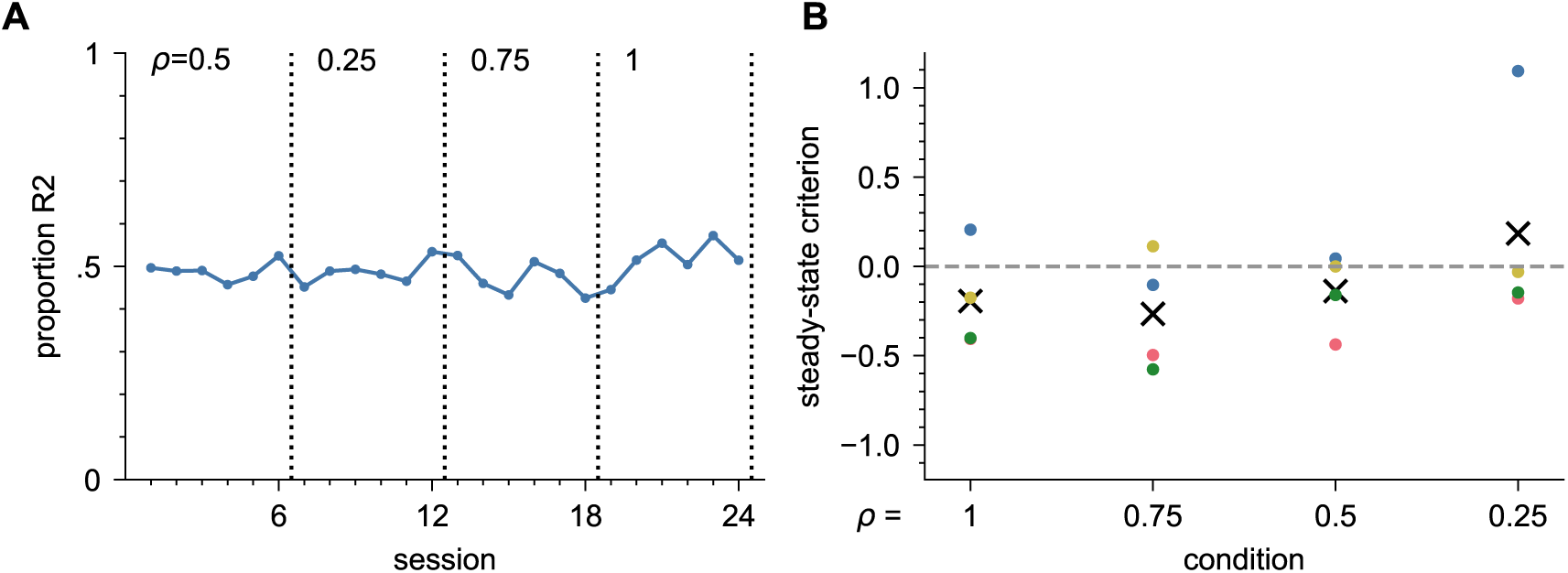
Behavior in Experiment 4 (reward density varied). **A)** Response proportions for example subject 9. Each data point is the proportion of R2 responses in one session. **B)** Steady-state criterion for all subjects. Each data point stems from one experimental condition, with colored dots representing individual subjects and means across animals shown as an ×. Conditions are ordered from high to low reward probability.

All conditions share the same equilibrium criterion within each of the models, too (*c_eq_* = 0 for the DT and RL model, *c_eq_* dependent on the model parameters for the modified KDB model). However, the criterion is still updated from trial to trial, and hence varies around the equilibrium. To investigate if the amount by which the criterion is updated depends on the reward density, we introduced a version of each model that allows different learning rates (i.e., Δ in the modified KDB model, Δ_max_ in the DT model, and *α* in the RL model, respectively) for conditions with different reward densities.

We fitted each of the models to the data from Experiment 4, both in its single-learning-rate version and the version with multiple learning rates (see Section 2.5.4 for details). The multiple-learning-rate models had one learning rate parameter for each condition. The fits and simulations of the single-learning-rate models for an example subject are shown in Figure 9; for the other subjects, they can be found in Figure E.1 in the appendix. The multiple-learning-rate models did not perform significantly better, so they were discarded, as we will describe next.

**Figure 9:**
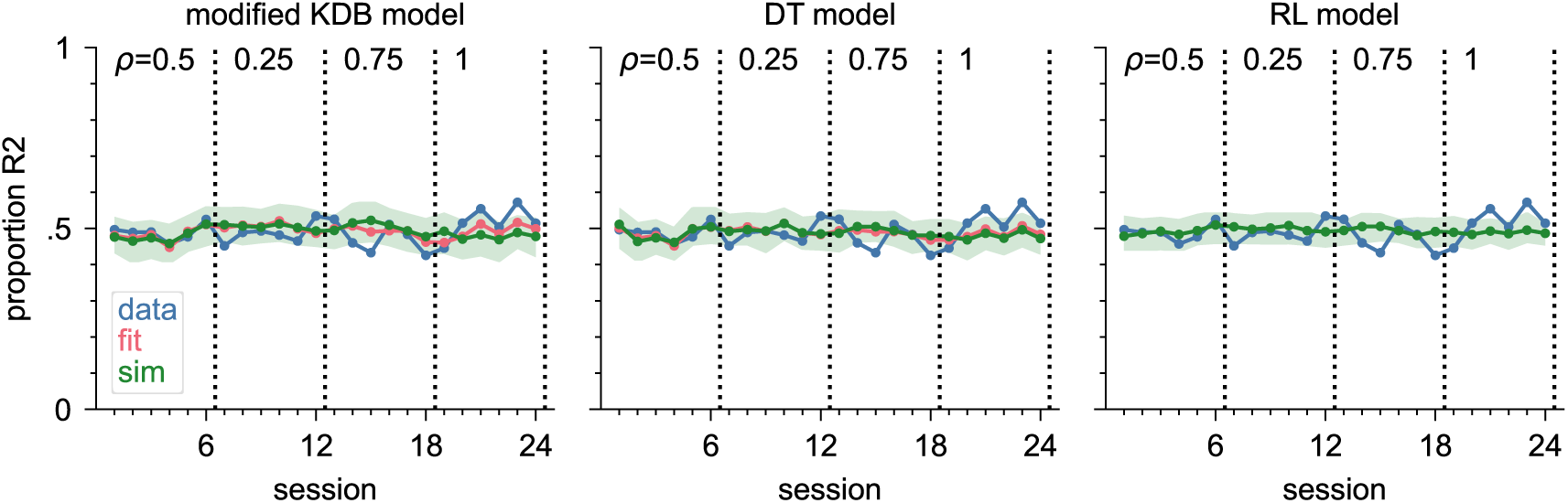
Model fit and simulations applied to the data of example subject 9 in Experiment 4. Each data point is the proportion of R2 in one session. Blue line: data. Red line: proportion of R2 predicted by the model fit (using the best-fitting parameters), i.e., session average of P(R2) in each trial given the actual trial history. Green line: median of 100 simulations (using the best-fitting parameters), shaded green area: central 95% of the simulations.

We compared the goodness-of-fit between the two versions for each model. Since the two versions have a different number of parameters, a direct comparison of the NLLs is not informative; so instead we computed the Bayesian information criterion (BIC, see Section 2.6 for details) and compared that between model versions. If the BIC is lower for the model version with multiple learning rate parameters, this justifies the inclusion of these additional parameters. The BIC comparison for Experiment 4 is shown in Table 6. The models with multiple learning rates consistently performed worse than the corresponding single-learning-rate models, with only a few exceptions. From a modeling perspective, this means that allowing a different learning rate for each condition did not improve the model and we can discard the multiple-learning-rate versions. Moreover, in the multiple-learning-rate versions the fitted values of learning rates in the different conditions showed a huge variation in their magnitude, but did not have any particular order (see Figure 10). This strengthens the conclusion that there was no systematic effect of reward density on the learning rate for any of the models. From a behavioral perspective, these findings indicate that reward density did not affect the animals’ learning rate.

**Figure 10:**
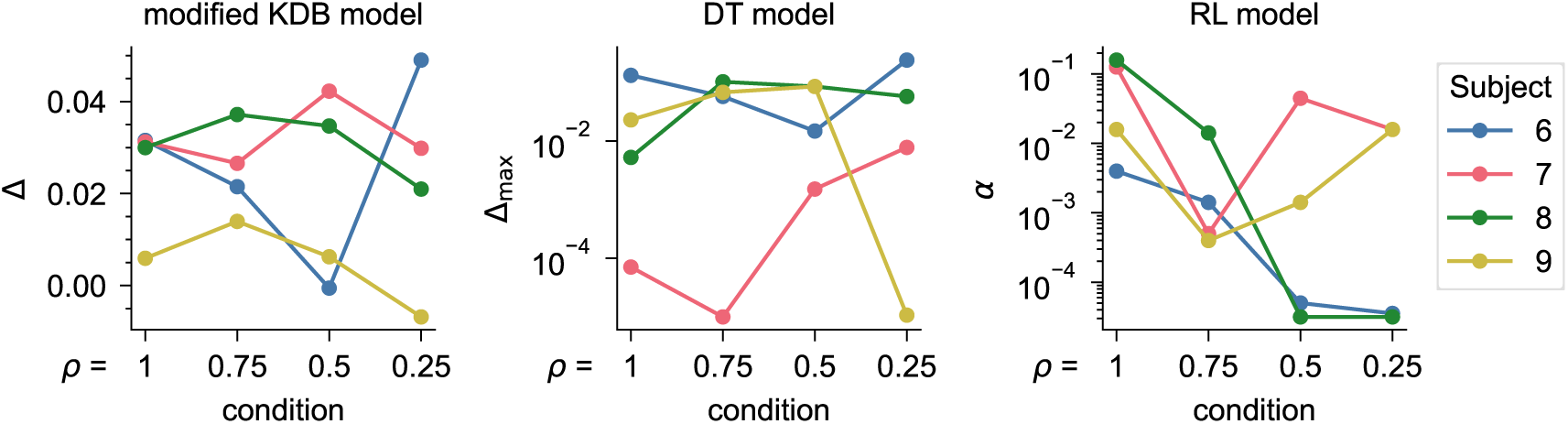
Best-fitting learning rate parameters in Experiment 4 for each model’s multiple-learning-rate version. The respective parameters are: modified KDB – Δ, DT – Δ_max_, RL – *α*. Note that the values for *α* are log-scaled. Conditions are ordered according to their reward probability. Each colored line represents results from one subject.

**Table 6:** BIC comparisons for Experiment 4. For each model, the single-learning-rate version is compared to the multiple-learning-rate version. The version that performs better (lower BIC) for each subject is colored green.

| subj. | modified KDB |  | DT |  | RL |  |
| --- | --- | --- | --- | --- | --- | --- |
|  | single | multi | single | multi | single | multi |
| 6 | 12634 | 12603 | 12619 | 12614 | 13380 | 13386 |
| 7 | 10714 | 10739 | 10742 | 10761 | 11336 | 11338 |
| 8 | 10188 | 10510 | 10515 | 10535 | 11382 | 11330 |
| 9 | 12181 | 12199 | 12177 | 12199 | 12181 | 12205 |

### 3.4 Experiment 5

While the model-based analyses of Experiment 4 suggest that reward density does not systematically affect learning rate, it could be that reward density does play a role when animals are additionally confronted with asymmetric RPs. In Experiment 5 therefore, RR and reward density were jointly varied while stimulus probabilities were again kept constant and symmetric. Besides a baseline condition with *ρ*_1_ = *ρ*_2_ = 0.5, reward ratios of 2:1 and 4:1 were tested, each at one higher and one lower reward density level and favoring each response option (see Table 1E). The behavior of an example subject is shown in Figure 11A and steady-state criteria in each condition for all subjects are shown in Figure 11B. As expected, there is a clear relation between RR and steady-state criterion, with more extreme reward ratios producing a stronger bias towards the response associated with the higher reward probability. This matches the observations from Experiment 2, where only RR was varied. In contrast, the steady-state criteria do not exhibit any effect of reward density. Within pairs of conditions that share the same ratio and differ only in reward density, there is no consistent pattern that would imply that higher reward density would lead to more or less extreme steady-state criteria.

**Figure 11:**
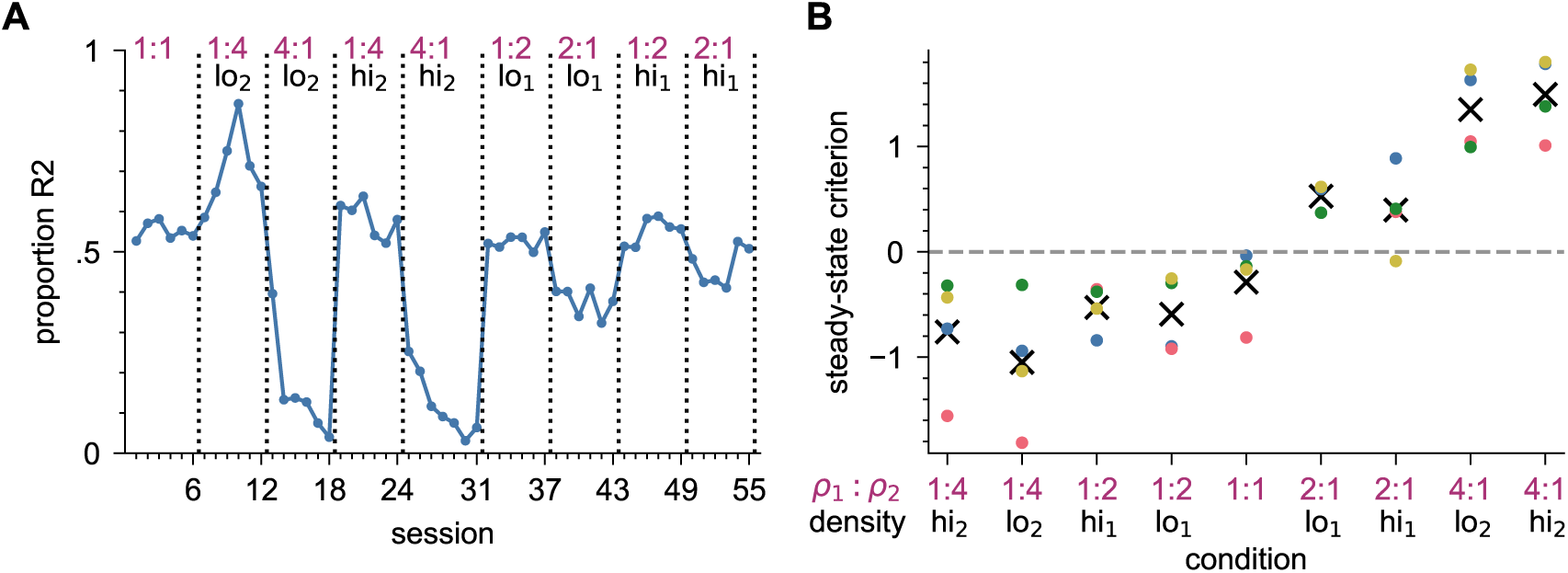
Behavior in Experiment 5 (RR and reward density varied) **A)** Response proportions for example subject 9. Each data point is the proportion of R2 in one session. **B)** Steady-state criterion for all subjects. Each data point stems from one experimental condition, with colored dots representing individual subjects and means across animals shown as an ×. Conditions are ordered from RP most strongly favoring response 2 towards most strongly favoring response 1, with high density conditions further towards the ends than the corresponding low density conditions.

A DT law analysis (see Figure E.5 in the appendix) shows that the DT law reasonably describes the relationship between response ratios and reward ratios in this setup, too, and reward density does not systematically affect this relationship.

To investigate if reward density influences the amount by which the decision criterion is updated in this setup, where reward density and RR are jointly varied, we fitted each of the models to the data from Experiment 5 in its single-learning-rate and a multiple-learning-rates version. In the multiple-learning-rates version, we grouped conditions with the same reward density together (e.g., conditions ‘2:1 hi_1_’ and ‘1:2 hi_1_’ into group ‘2 hi_1_’) and fitted the model with one learning rate for each condition group. The fits and simulations for an example subject are shown in Figure 12; for the other subjects they can be found in Figure E.2–E.4 in the appendix (one figure per model).

**Figure 12:**
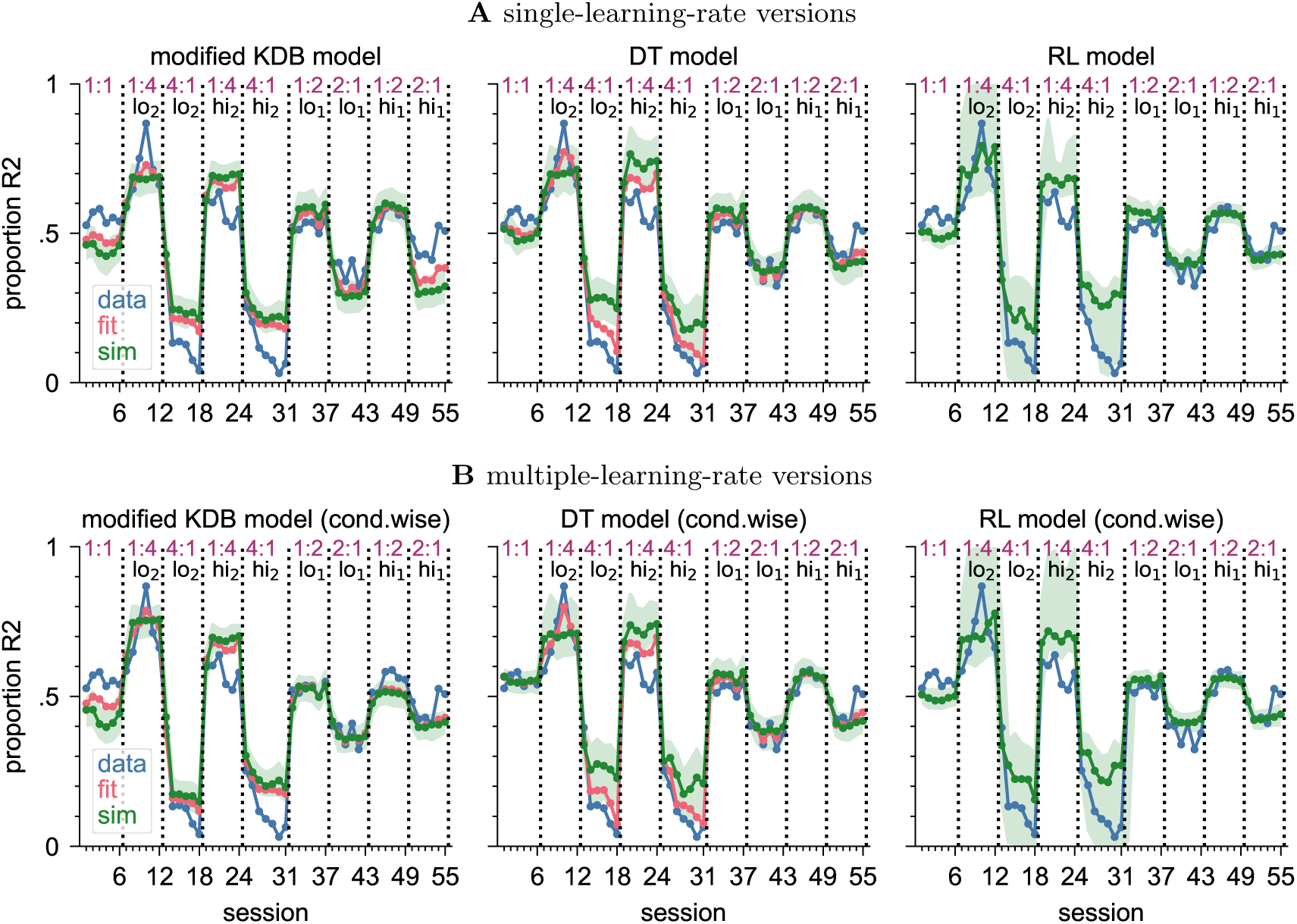
Model fit and simulations applied to the data of example subject 9 in Experiment 5. **A)** Single-learning-rate model versions. **B)** Multiple-learning-rate model versions. Each data point is the proportion of R2 responses in one session. Blue line: data. Red line: proportion of R2 predicted by the model fit (using the best-fitting parameters), i.e., session average of P(R2) in each trial given the actual trial history. Green line: median of 100 simulations (using the best-fitting parameters), shaded green area: central 95% of the simulations.

In Experiment 5 – in contrast to Experiment 4 – the steady-state criterion also varies between conditions (as seen in Figure 11B), since reward ratios are varied. Therefore, reward-density dependent learning rates would not only manifest in differences in trial-to-trial criterion updating, but would also on a macro-level lead to different speeds of adaptation to the new steady state in the different conditions.

We again compared the goodness-of-fit between the two versions for each model using the BIC (cp. Section 2.6). The results are shown in Table 7. Here, the models with multiple learning rates always performed better than the corresponding single-learning-rate models. However, the fitted values still varied unsystematically. Figure 13 shows the fitted learning rates for each subject for each condition group. Comparing the respective high-density and low-density condition groups to each other (i.e. 2 hi_1_ vs. 2 lo_1_, 4 hi_2_ vs. 4 lo_2_), no observable pattern emerged. For each model, sometimes the higher densities corresponded to higher learning rates and sometimes to lower learning rates, both within and across subjects. Since this absence of a relation seemed surprising in the face of the BIC comparisons that indicate that the variations in learning rate between conditions with different reward densities are meaningful, we fitted another model version with one learning rate for each individual condition. This further improved the BIC in comparison to the models with one learning rate per reward density. Therefore, while there seem to be meaningful variations in learning rate between conditions, this is apparently not an effect of reward density.

**Figure 13:**
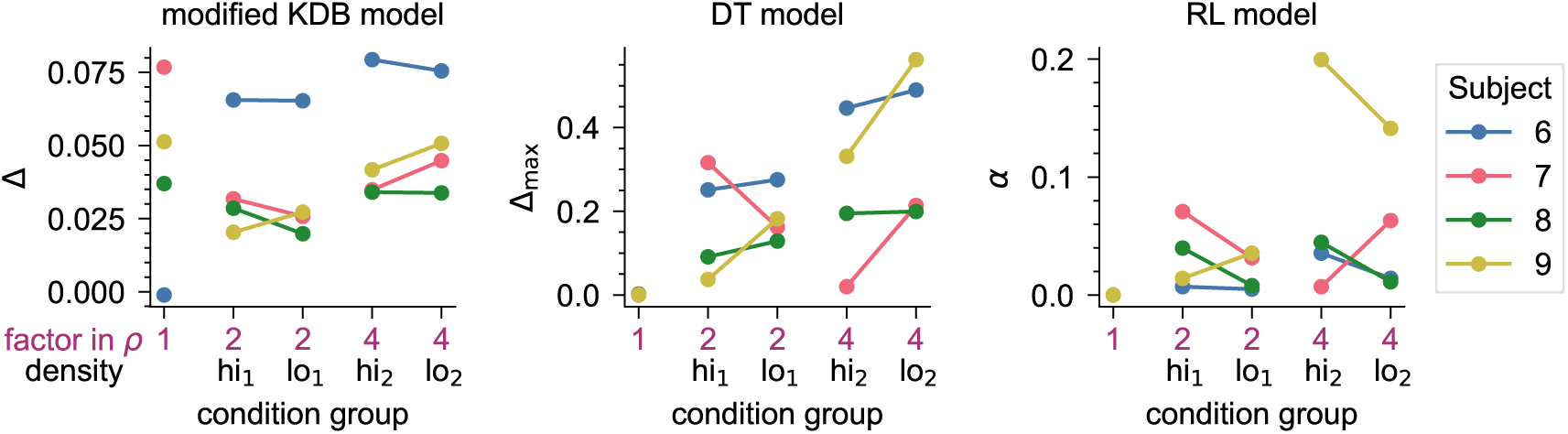
Best-fitting learning rate parameters in Experiment 5 for each model’s multiple-learning-rate version. The respective parameters are: modified KDB – Δ, DT – Δ_max_, RL – *α*. Conditions with the same factor between the two reward probabilities were grouped together (e.g. *ρ*_1_:*ρ*_2_ = 2:1 and *ρ*_1_:*ρ*_2_ = 1:2 → factor 2). Each colored line represents results from one subject.

**Table 7:**
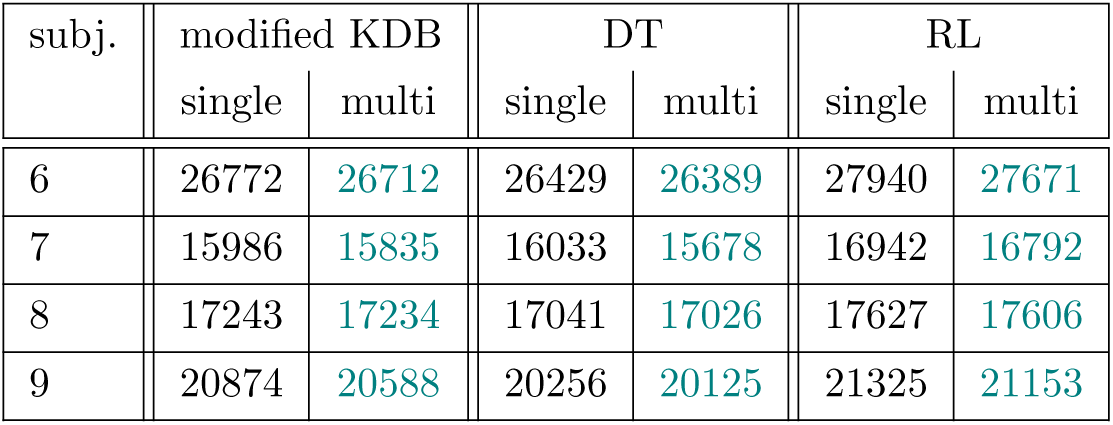
BIC comparisons for Experiment 5. For each model, the single-learning-rate version is compared to the multiple-learning-rate version. The version that performs better (lower BIC) for each subject is colored green.

| subj. | modified KDB |  | DT |  | RL |  |
| --- | --- | --- | --- | --- | --- | --- |
|  | single | multi | single | multi | single | multi |
| 6 | 26772 | 26712 | 26429 | 26389 | 27940 | 27671 |
| 7 | 15986 | 15835 | 16033 | 15678 | 16942 | 16792 |
| 8 | 17243 | 17234 | 17041 | 17026 | 17627 | 17606 |
| 9 | 20874 | 20588 | 20256 | 20125 | 21325 | 21153 |

## 4 Discussion

We conducted five experiments to characterize the effects of SPRs, RRs, and reward density on criterion learning, both at the steady-state and the trial-by-trial level. Our main findings are:

First, while SPR and RR manipulations similarly biased choices towards the more probable (SPR) and more profitable (RR) response options, the effect of RR on criterion setting was consistently larger than that of SPR (Experiments 1 and 2). Moreover, the effect of RRs was clearly more pronounced than that of SPRs in Experiment 3 which directly compared the influence of these two factors. Relatedly, the three trial-by-trial models provided reasonable fits to the data of Experiments 1 and 2 (except for the RL model in Experiment 1), but fitted learning rates were more than an order of magnitude larger for Experiment 2, i.e., when RRs were manipulated. All models failed at generating behavior like the experimental subjects in Experiment 3 in which SPRs and RRs were concomitantly manipulated.

Second, changes in reward density did not systematically affect learning rates (Experiment 4 and 5). When reward density and RR were manipulated concomitantly, equipping the models with different learning rates for different conditions improved their fit, despite the lack of relation between learning rates and reward density (Experiment 5).

### 4.1 Differential effects of SPR and RR manipulations on criterion setting

The effect of RR manipulations on criterion setting was consistently larger than that of SPR manipulations (Figure 3C). This difference is also reflected in some of the fitted model parameters (larger *a* in the DT model, and a larger ratio 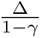 in the modified KDB model), which led to more extreme steady-state criteria in Experiment 2 (RR) than in Experiment 1 (SPR). Moreover, the fitted learning rates of the three models differed between the two manipulations: they were substantially larger in Experiment 2 (RR) than in Experiment 1 (SPR; Tables 3, 4 and 5).

It seems likely that the differential effects of SPR and RR are also responsible for the bad fits observed for the choice data in Experiment 3. In this experiment, we explicitly juxtaposed SPRs and RRs of the same magnitude to compare their effect sizes. Importantly, an ideal observer within the SDT framework should consistently opt for a neutral criterion (i.e., a criterion value of 0) in all experimental conditions. Moreover, the DT and modified KDB models each predict the same steady-state criterion for all conditions in this experiment. Nonetheless, animals were consistently biased towards the choice option with the higher reward probability (Figure 6BC). Together, these three experiments clearly demonstrate that SPR and RR modifications do not affect criterion learning equally. Therefore, any model that does not account for this finding, is incomplete or wrong.

While SPR and RR equally determine the optimal decision-theoretic criterion (s.a.), an animal subjected to these manipulations will experience very different stimulus-response-outcome contingencies. For illustration, assume a subject transitions from an RR schedule of 1:1 to 4:1, while maintaining a neutral decision criterion of *c* = 0. When the RR changes, the animal will experience a change in response-contingent reward probabilities (i.e., increased *P* (Reward|*R*1) and decreased *P* (Reward|*R*2)), and a larger fraction of all rewards will be obtained from R1. However, when the SPR schedule changes from 1:1 to 4:1, the *P* (Reward|*R*1) and *P* (Reward|*R*2) stay the same. Instead, stimulus 1 probability *π*_1_ is four times as high as *π*_2_, and in consequence, the response proportion P(R1) will be four times higher than P(R2) as well, which then results in a larger fraction of all rewards coming from R1 (i.e., (*P* (Reward|*R*1) ∗ *P* (*R*1))*/*(*P* (Reward|*R*2) ∗ *P* (*R*2)) = 4). This line of reasoning illustrates an important point: we need to consider which of the above (and related) probabilities the animals keep track of, and how. From the differential effects of the SPR and RR manipulations, we hold it likely that criterion learning is not solely influenced by stimulus-independent, response-contingent reward rates, as implied by the KDB and DT models. Instead, we suggest that the influence of a reward is determined not only by the response which produced it, but also depends on the stimulus evidence. Indeed, the RL model follows this idea, as it updates action values based on an assessment of the current stimulus. However, it does not keep track of the stimulus history, which could be a reason that the RL model still failed to account for the animals’ response patterns.

The RL model’s behavior depends on some assumed prior stimulus distributions. We chose these distributions to match the real stimulus distributions with *π*_1_ = *π*_2_ = 0.5, reflecting the animals’ extensive experience with the stimuli before the first experiment began. This allows the model to fit the data in Experiment 2, which features such a scenario. However, the original model by Lak et al. (2020b) was not designed to work in a scenario with changing SPPs, like Experiments 1 and 3, and our choice of stimulus prior does not address this issue, as it still assumes a fixed overall stimulus distribution. Simply using a differently (correctly) weighted version of the two stimulus distributions in each condition would not be plausible, because the animals cannot know about that weighting at the outset of a condition. Thus, the RL model would need an additional mechanism to learn or update the prior distribution in order to be a suitable model for behavior in experiments where the stimulus distributions do not stay fixed. Indeed, Fritsche et al. (2026) have recently proposed a model that represents stimulus priors and updates them continuously through sensory prediction errors. Moreover, they showed that striatal dopamine activity differs between SPR and reward magnitude conditions, although behavioral bias was very similar. It remains to be seen whether a model representing stimulus priors only can adequately capture our data, or whether a successful model requires the explicit representation of stimulus distributions. The latter may become especially relevant when non-Gaussian stimulus distributions are used (Menichini et al., 2025). For studies in humans it even seems desirable to describe mechanisms that can adapt to arbitrary, non-fixed stimulus distributions (Turner et al., 2011).

Previous authors have also described differential effects of SPR and RR. Most notably, McCarthy and Davison (1979) asked whether the established biasing effects of SPRs arise indirectly through the effect of asymmetric reward frequencies. They “clamped” relative RRs in a so-called controlled reinforcer ratio (CRR) procedure in which the RR was independent of changes in SPRs and found that SPR changes by themselves did not affect response bias in this scenario. They concluded that SPR is not an effective biasing variable by itself, but biases choices only indirectly through the different RRs which result from such manipulations in non-CRR procedures. Our findings are in opposition to their claim that the biasing effect of SPRs is mediated exclusively by unequal reward rates for the two choices: if this were true, then we would expect that that SPR and RR manipulations would lead to similar parameter values in Experiments 1 and 2. Moreover, SPRs and RRs should exert the same effect in Experiment 3. This was clearly not the case. Notably, McCarthy and Davison (1979) already observed that SPR and RR manipulations lead to different sensitivities to reward.

Interestingly, in humans SPR manipulations generally produce larger (and quasi-optimal) criterion shifts than manipulations of RRs. In a series of studies, Maddox and coworkers (for review, see Maddox, 2002) documented that (steady-state) criteria with SPR manipulations are very close to optimal (accuracy-maximizing) values, while criteria under RR manipulations were consistently less extreme than optimal. However, we found just the opposite – SPR manipulations yielded less extreme criterion values. Importantly, Maddox and coworkers have always manipulated RRs by changing reward magnitudes, with reward probability being 1 throughout. The effect of changing reward probabilities have, to our knowledge, not been investigated in humans. Although these two manipulations might appear similar, they could very well instantiate different behaviors. For example, when reward magnitude is manipulated, subjects receive feedback in each and every trial. In our experiments, unrewarded and incorrect responses yielded the same outcome (a short time-out), so feedback was unambiguous only when reward was provided. It is thus entirely possible that our results had been different if we had operationalized RRs through reward magnitude rather than reward probability ratios.

The divergent effects of SPRs and RRs are not well captured by any of our models. The simplest explanation for this is that none of the models explicitly represents stimulus probabilities. Therefore, adding a malleable representation of stimulus probabilities (or at least their ratio) might render the models suitable to fit the data of Experiments 1 through 3.

### 4.2 Reward density has no consistent effect on criterion learning

In Experiment 4, we varied overall reward density, keeping SPRs and RRs constant at 1:1. We fitted the animals’ choice data with models featuring either a single learning rate or multiple learning rates (one for each reward density). We hypothesized that learning rates depend on reward density, such that low-reward schedules would be associated with higher learning rates. Instead, we observed no consistent effect of reward density manipulations (Figure 10). On the one hand, this finding is consistent with the matching law and, by extension, the DT law, which both posit that response bias is a function of the reward ratio, which is by definition independent of reward density. Also, this lack of effect is predicted by both the modified (but not the original) KDB model as well as the DT model. On the other hand, effects of reward density on behavior have been found elsewhere. For example, monkeys exhibit a win-stay strategy in high-profit environments but a lose-switch strategy in low-reward environments (Wittmann et al., 2020), and behavior in a certain environment depends on whether that environment is entered following exposure to a high-profit or low-profit reward schedule, a presumably adaptive strategy (McNamara et al., 2013). However, in these studies, subjects were not confronted with perceptual uncertainty.

Experiment 4 manipulated reward density alone to isolate its effects. In Experiment 5, we manipulated reward density in combination with reward ratios, the rationale being that Experiment 4 might fail to show an effect because little learning is required when neither SPR nor RR changes. Indeed, for each of the three models, allowing multiple learning rates considerably improved the goodness of fit for all subjects when controlling for the larger number of free parameters (Table 7). That said, there was no systematic trend for learning rates to be higher or lower in low-reward environments, and this was true regardless of the chosen model (Figure 13). This indicates that there is some other aspect of the conditions, i.e., the interplay between RR and reward density, that none of the three models captures. A different learning rate for each condition allows the models to fit the learning trajectory and steady-state in each condition independently and thereby improves the fit, but an explanation which aspects are relevant and in what way they determine the parameters for each condition is lacking.

### 4.3 Conclusions and future directions

Despite their surface similarity, SPR and RR manipulations exert different effects on the animals’ behavior. In a similar vein, one might ask whether the effects of reward probability and reward magnitude manipulations are different; to our knowledge, this question has not been investigated so far. In conclusion, every successful trial-by-trial model of criterion learning needs to account for the differential effects of SPR and RR. To that end, it will be necessary to investigate in future experiments whether animal subjects keep track of stimulus priors, stimulus distributions, or both, and to investigate how stimulus learning can best be accommodated into existing models.

## Acknowledgements

Preparation of this work was supported by grants from the Deutsche Forschungsgemeinschaft to M.C.S. (project IDs 424828846 and 543128862) and F.J. (project ID 424828846). The funders had no role in study design, data collection and analysis, decision to publish, or preparation of the manuscript.

## A Equilibrium criterion of the modified KDB model

An equilibrium is reached when the expected change in criterion from trial to trial is 0, i.e., 𝔼[*c*(*t*+1)] = *c*(*t*). Let *P*_1_ = *P* (*R*1*, Rw* = 1) denote the probability that R1 is emitted and rewarded in a trial and *P*_2_ analogously for R2. Using the criterion update formulas for the modified KDB model (Equations 9, 10 and 11), it thus holds that

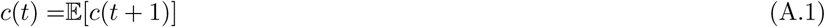

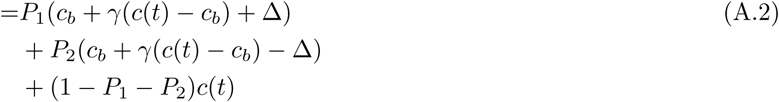

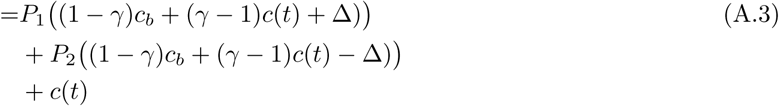

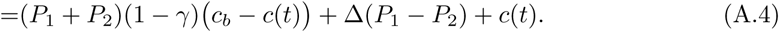

This is equivalent to

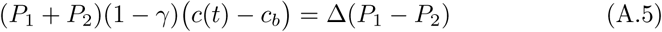

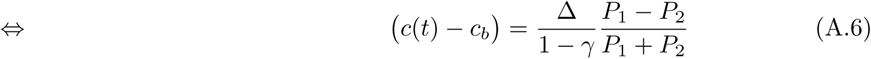

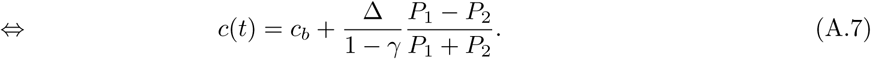

From this it can be seen that the ratio 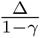 scales the equilibrium criterion, i.e., the two parameters Δ and *γ* jointly influence its position. The higher the ratio, the more extreme the equilibrium criterion will be, for any given experimental condition. An example visualization is shown in Figure 2A.

Since *P*_1_ = *P* (*R*1|*S*1)*π*_1_*ρ*_1_ and *P*_2_ analogously, the difference in relative reward probabilities, (*P*_1_ − *P*_2_)*/*(*P*_1_ + *P*_2_), can be expressed as

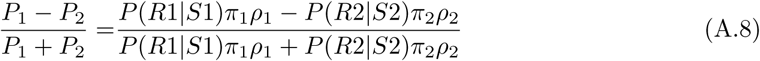

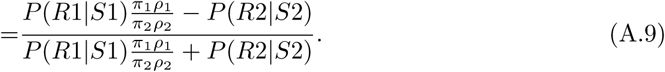

This shows that the equilibrium criterion depends on the experimental condition, and is influenced by SPP and RP equally, as it is a function of 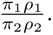 If no rewards were given, *P*_1_ = *P*_2_ = 0 and the equilibrium criterion would be the subject’s inherent bias criterion *c_b_*.

The response probabilities *P* (*R_i_*|*S_i_*) themselves depend on *c*(*t*), therefore, to determine the equilibrium criterion, Equation A.7 has to be solved for *c*(*t*). This cannot be done analytically, so numerical methods have to be used.

## B Fitting the multiple-learning-rate versions of the models

The multiple-learning-rate versions of the models have one learning rate parameter *LR_g_*(Δ*_g_* in the modified KDB model, 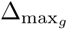 in the DT model, and *α_g_*in the RL model) for each trial group *g*.

For the modified KDB model, the multiple-learning-rate can be fitted to the data in the same way as the single-learning-rate version. Even with a different Δ for each trial group, the log-likelihood for a fixed *γ* can still be expressed as a generalized linear model of all other parameters and hence be fitted using standard numerical optimization methods.

For the DT model, the multiple-learning-rate version was fitted to the data as follows. Starting with the first trial group in the experiment, here denoted as *g*1, the log-likelihood for each parameter combination in a grid of 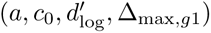 was computed for data from the trials in this group only. For each combination of 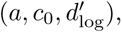 the best Δ_max*,g*1_ is the one that leads to the highest log-likelihood of the data in *g*1. Then, for each combination of 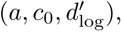 the log-likelihood of the data from the trials in group *g*2 was determined for each Δ_max*,g*2_ and the best one chosen.^8^ The same procedure was executed for all trial groups consecutively, overall yielding for each parameter combination 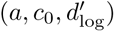 the best Δ_max*,g*_ as well as the corresponding log-likelihood of the data in each trial group *g*. The total log-likelihood for the full data set was then computed by summing up the log-likelihoods of all trial groups. The values of 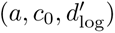 for which the total log-likelihood was highest were chosen as the best-fitting parameters. Note that this method requires computing the log-likelihoods for the full range of candidate values for 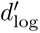 instead of selecting the best one in each trial group, since the best 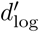 in a trial group is not necessarily the best one overall. Therefore, 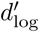 cannot be determined by fitting a generalized linear model as described in Section 2.5.2, but another grid dimension has to be added. The range of 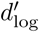 used for the grid was 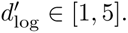

For the multiple-learning-rate version of the RL model, the fitting procedure worked similarly. Here, the initial values *V*_1_(0) and *V*_2_(0) are not free parameters but fixed to 0.5. Starting with the first trial group *g*1, the NLL_alt_ for each parameter combination on the grid *σ*^2^*, α_g_*_1_ was computed for data from the trials in trial group *g*1. For each *σ*^2^, the best *α_g_*_1_ was chosen and used to compute *V*_1_ and *V*_2_ at the end of trial group *g*1, i.e., at the beginning of group *g*2. The same procedure was then applied sequentially to all trial groups, for each *σ*^2^ yielding the best {*α_gi_*} for all groups *g_i_*.

As for the DT model, the total NLL_alt_ was then computed by summing up the NLL_alt_ of all trial groups, and the value for *σ*^2^ with the corresponding best {*α_gi_*} was chosen that led to the highest total NLL_alt_. The grid range for all *α_gi_* was chosen as for the single-learning-rate version of the model. For *σ*^2^, the range was extended to go up to 1, as some of the found maxima were located at the edge of the smaller range.

## C Additional results for Experiment 1 & 2

### C.1 Model fits and simulations for all subjects

Figures C.1–C.3 and C.4–C.6 show the fits and simulations for all three models to the data from Experiment 1 and 2, respectively.

### C.2 DT law fits to model simulations

To see if the model simulations follow the DT law like the data does, we looked at the response ratios and reward ratios per stimulus for each simulation and fitted the DT law for each simulation. Figure C.7 shows the resulting central 95% of points (green ellipses) and central 95% of the fitted lines (shaded green area) for an example subject (all models) and Figures C.8–C.10 for all other subjects (one figure per model). In addition, we averaged each point over simulations and fitted the DT law to this simulation mean for each model, shown in the same figures as solid green line.

For the modified KDB and DT model (Figure C.7, left and middle column), comparing each real data point to the distribution of simulated points from the same condition again reveals that the variance in the original data is higher than the one exhibited by the model. Note that the actual variance of the model behavior is also higher than implied by the simulation data points though, because in a few cases the model exhibited exclusive choice behavior in one or several of the conditions. These conditions had to be excluded from the analysis, since exclusive choice means a response and reward ratio of ∞ or −∞. Therefore, the shown variability of the simulations is necessarily an underestimation of the real simulation variability. The straight lines fitted to the original data mostly fall within the central 95% of the straight lines fitted to the individual simulations, which indicates that the DT model can produce behavior that follows regularities that are similar to the ones of the animals’ behavior in the experiments.

For the RL model (Figure C.7, right column), the fitted straight lines for the simulation mean in Experiment 2 (RR varied) again agree relatively well with the fitted straight lines for the original data: The line for the data falls into or close to the central 95% of the straight lines fitted to the individual simulations. For Experiment 1 (SPR varied), however, the slopes of the lines fitted to the simulations (i.e., the sensitivity to reward *a*) are substantially lower than those of the lines fitted to the original data. This reinforces the impression that the RL model cannot accurately reproduce the behavior observed in the data for the setup with varying stimulus probabilities that was used in Experiment 1.

**Figure C.1:**
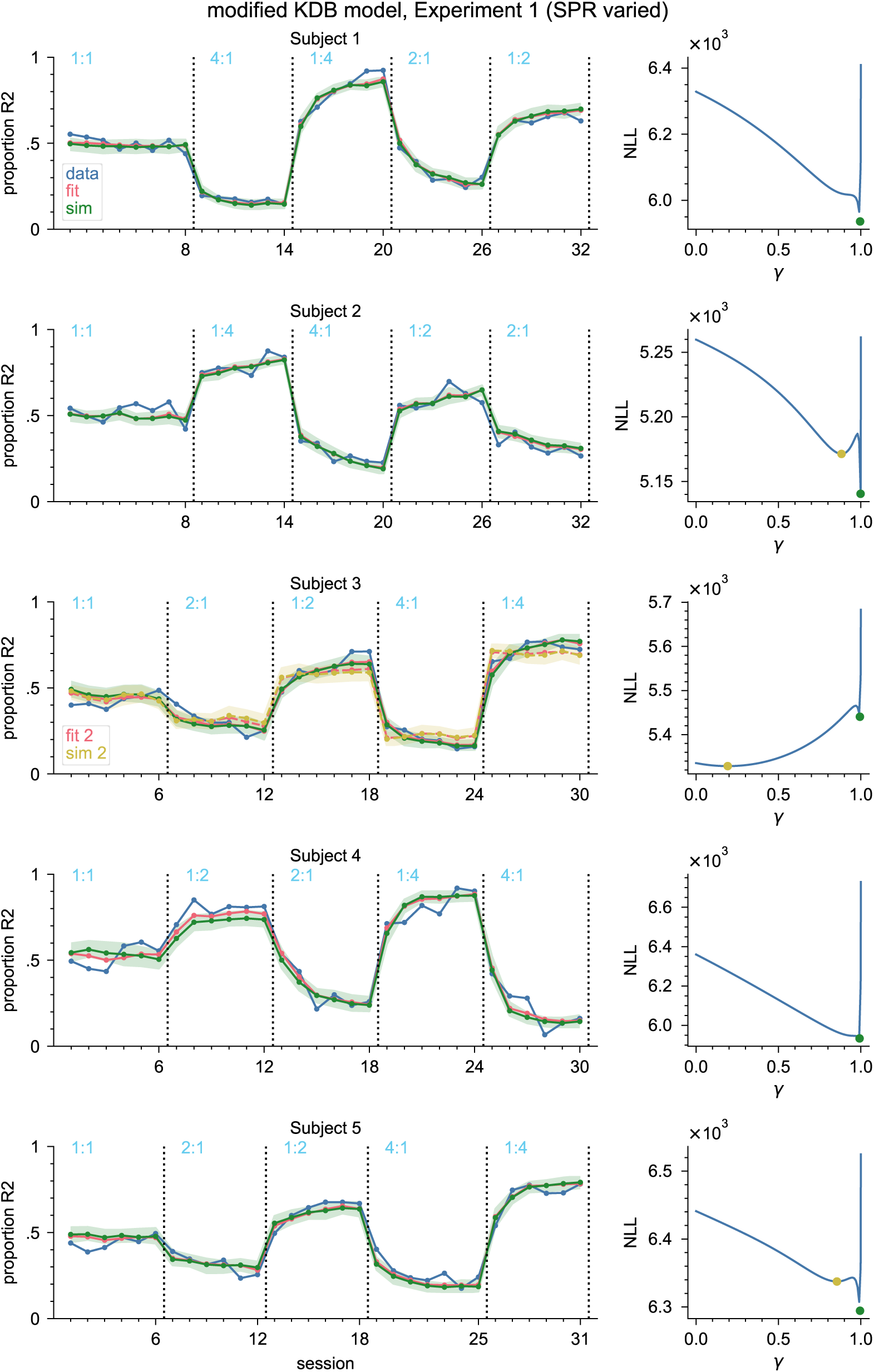
Results from fitting the modified KDB model to the data from Experiment 1 (SPR varied). Left: Model fit and simulations. See caption of Figure 5 for a detailed description. Wherever a second minimum exists in the NLL, the corresponding simulations are shown in yellow. Right: Best NLL for each value of *γ*. The minimum is marked in green; the second minimum - if one exists - is marked in yellow.

**Figure C.2:**
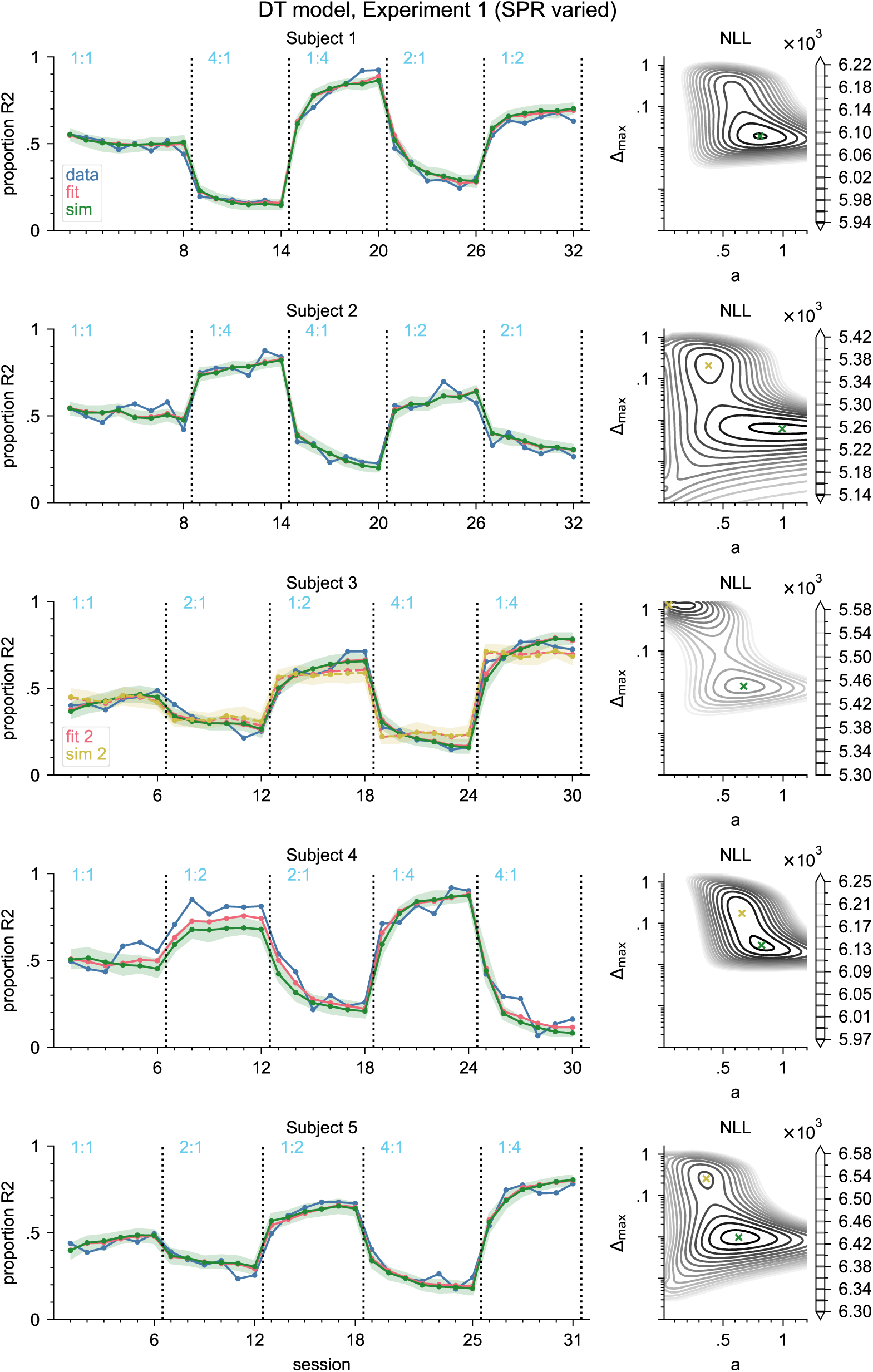
Results from fitting the DT model to the data from Experiment 1 (SPR varied). Left: Model fit and simulations. See caption of Figure 5 for a detailed description. Wherever a second minimum exists in the NLL, the corresponding simulations are shown in yellow. Right: NLL surface. The minimum is marked in green; the second minimum - if one exists - is marked in yellow. The y-axis is log-scaled for better visibility of both minima.

**Figure C.3:**
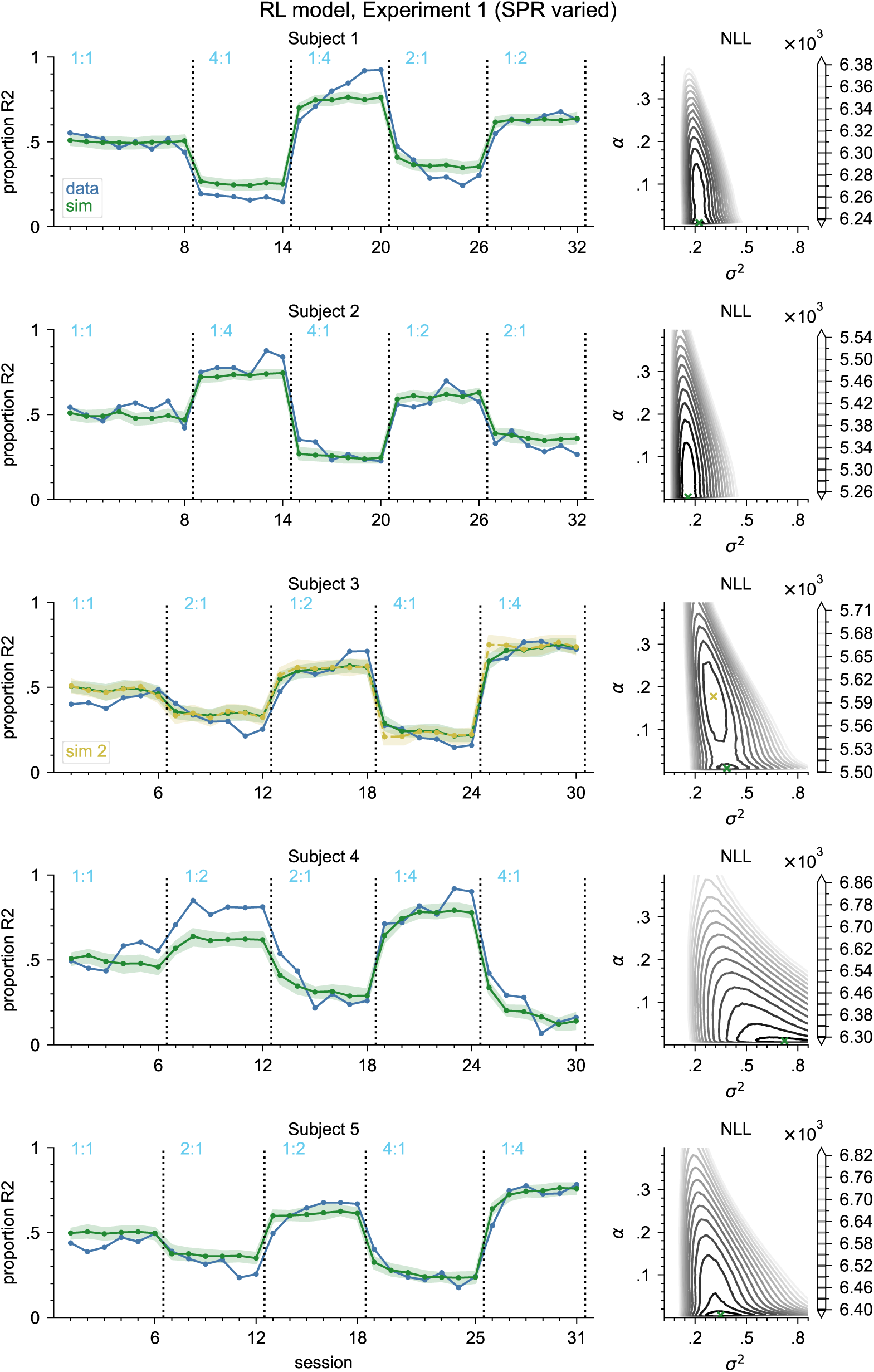
Results from fitting the RL model to the data from Experiment 1 (SPR varied). Left: Model fit and simulations. See caption of Figure 5 for a detailed description. Wherever a second minimum exists in the NLL_alt_, the corresponding simulations are shown in yellow. Right: NLL_alt_ surface. The minimum is marked in green; the second minimum - if one exists - is marked in yellow.

**Figure C.4:**
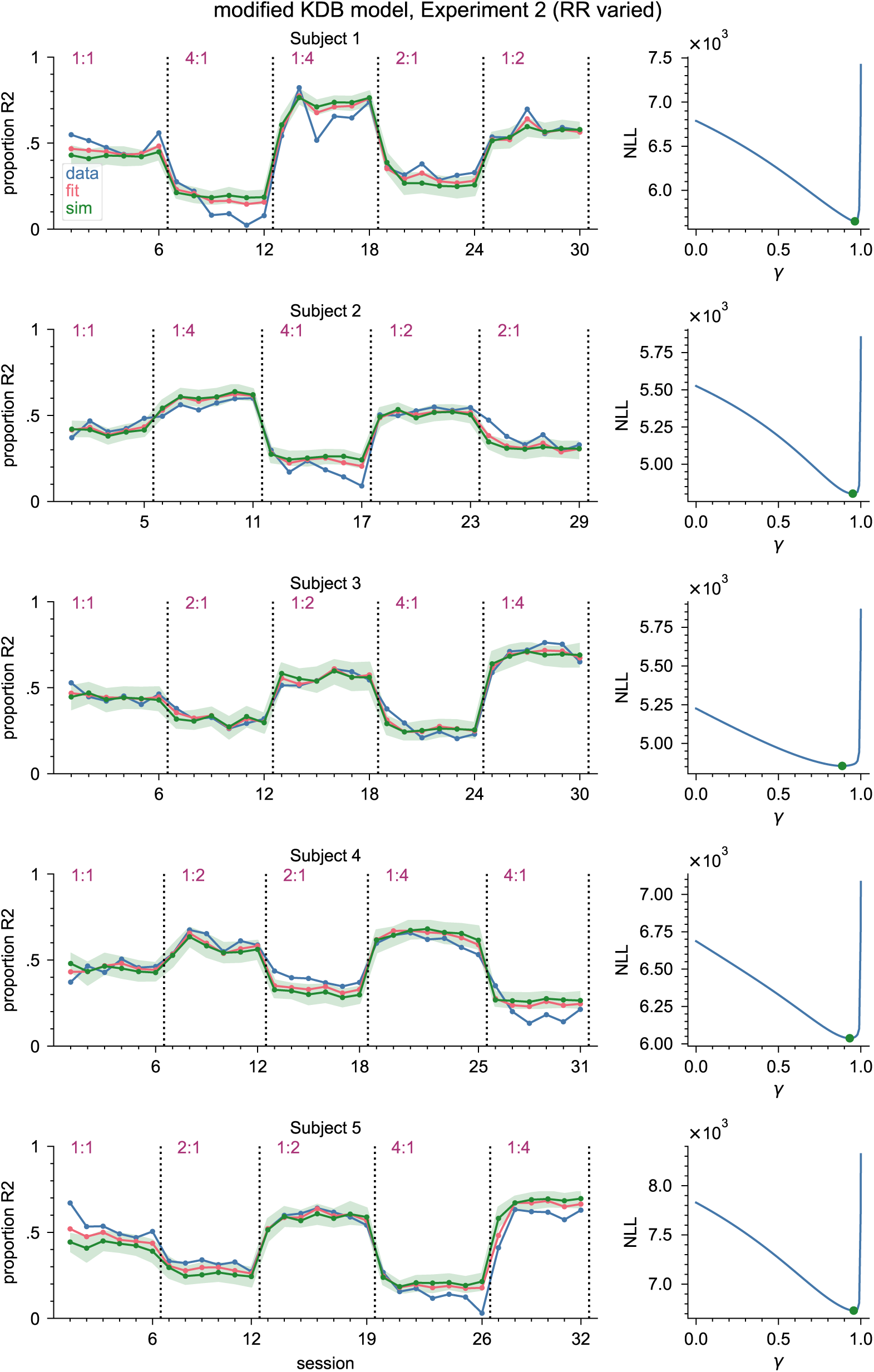
Results from fitting the modified KDB model to the data from Experiment 2 (RR varied). Same as Figure C.1, but for Experiment 2.

**Figure C.5:**
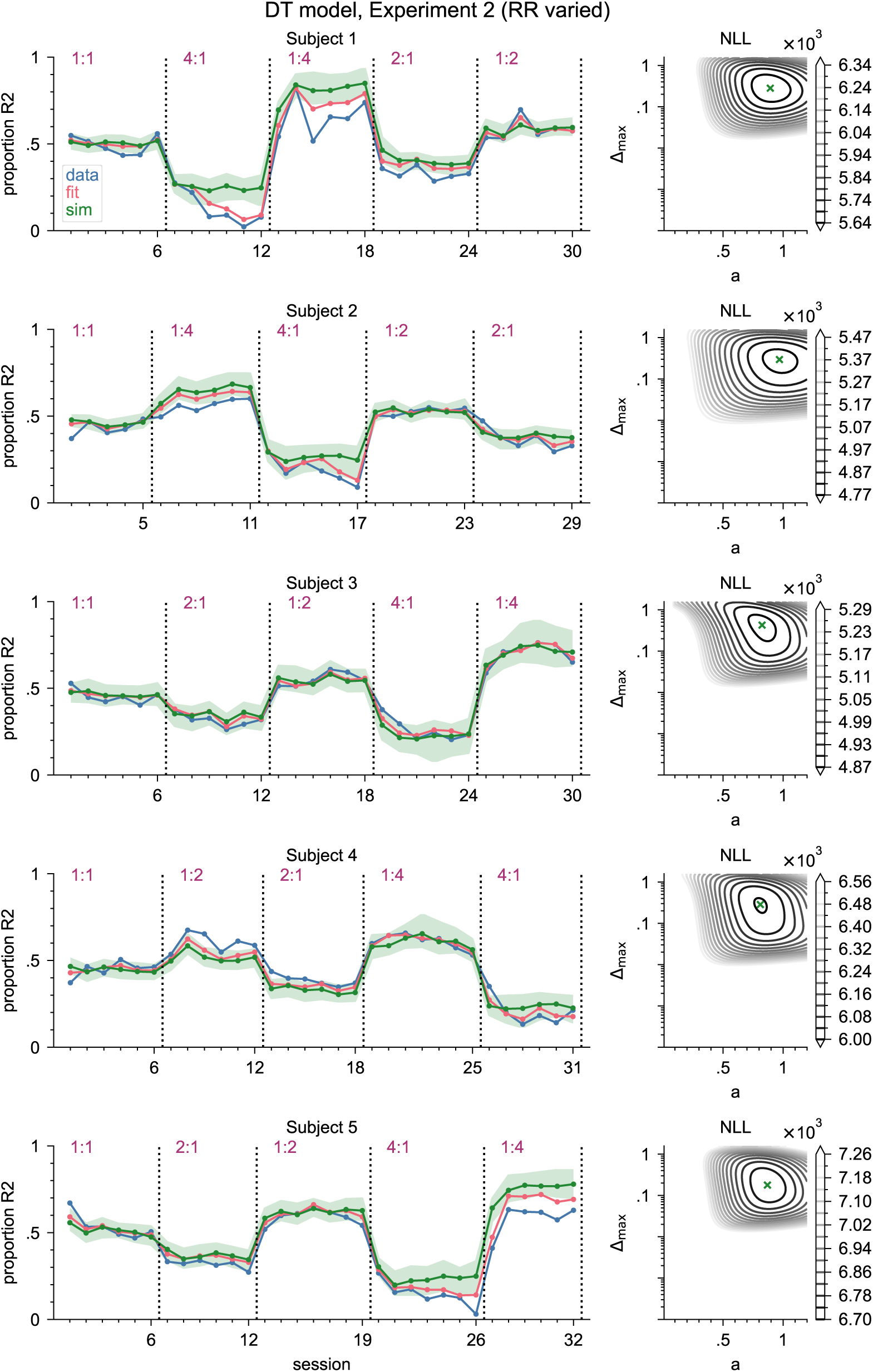
Results from fitting the DT model to the data from Experiment 2 (RR varied). Same as Figure C.2, but for Experiment 2.

**Figure C.6:**
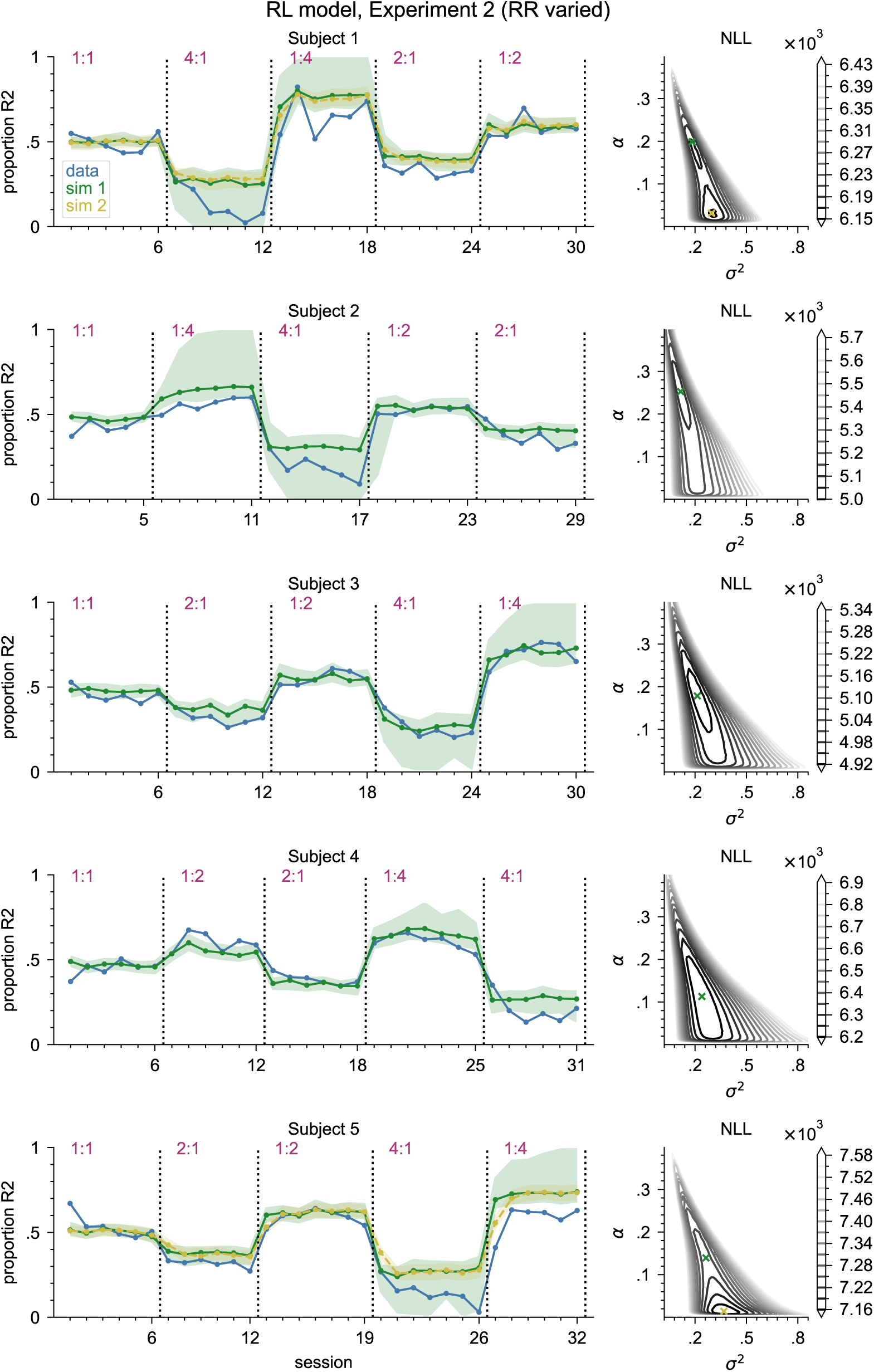
Results from fitting the RL model to the data from Experiment 2 (RR varied). Same as Figure C.3, but for Experiment 2.

**Figure C.7:**
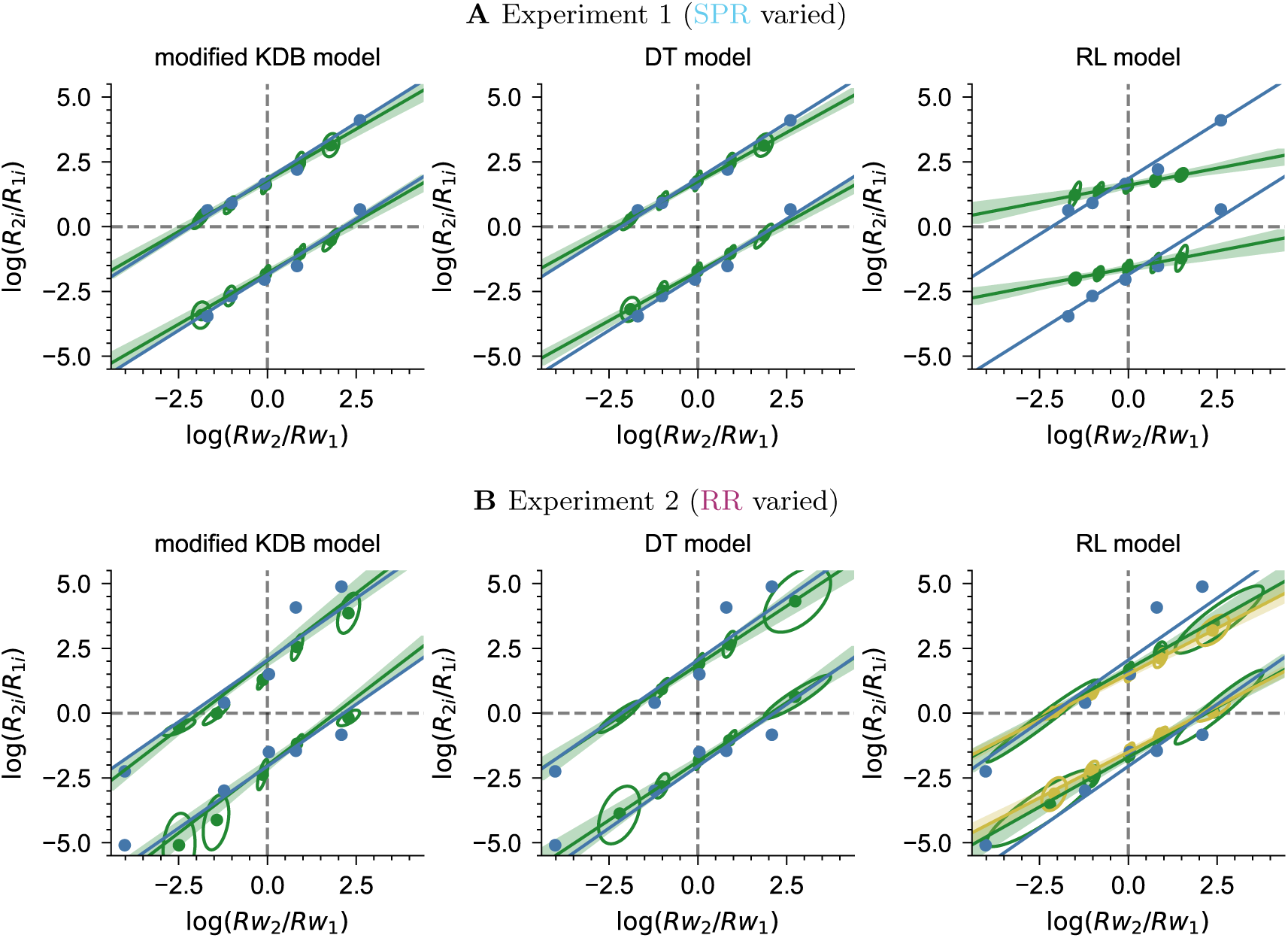
DT law fits for example subject 1 in Experiment 1 and 2. The DT law was fitted to the data (blue) as well as to 100 simulations from the model with the best-fitting parameters (green). Green dots and lines correspond to the DT law fit to the median of the simulations the shaded area marks the central 95% of the lines fitted to each simulation (at each point of the x-axis). The ellipses indicate the central 95% of simulations at each data point (assuming 2D gaussian distributions). Yellow: same as green, but for the parameters at the NLL minimum, in case these differ from the best-fitting parameters (cp. Table 5).

## D Additional results for Experiment 3

### D.1 Criterion development for all subjects

The criterion development for example subject 1 was shown in Figure 6B; for all other subjects it can be seen in Figure D.1.

### D.2 Model fits for Experiment 3

Figure 7 showed the models’ fits and simulation behavior for the data from Experiment 3 for example subject 1. For the other subjects, they are shown in Figure D.2.

It can be seen clearly that none of the models can produce behavior similar to the data.

What might seem surprising at first glance is that the modified KDB and DT model’s behavior also does not always align with the predictions resulting from their equilibrium criterion (a constant *c_eq_* as determined by Equation A.7 for the modified KDB model and *c_eq_* = 0 for the DT model throughout the whole experiment), marked as solid black lines. The simulations for these models often stay close to this equilibrium (in these cases, the median line is close to the reference line), but sometimes also diverge from that, approaching proportions of R2 close or equal to 0 or 1 (see shaded 95% interval, which extends downwards to or close to 0). That is because the models with the respective best-fitting parameters often have several stable equilibria: one that is close to *c* = *c_b_* (modified KDB) or *c* = 0 (DT), and one towards each side that is relatively far out. For subject 4, there is no stable middle equilibrium for the modified KDB model at all, but only an unstable one instead. As soon as the criterion by chance leaves the region of attraction of the middle equilibrium, which for the best-fitting parameters is often very small (or even non-existent), it quickly moves towards one of the other equilibria with almost exclusive choice.

Such an unstable behavior could be taken as an indicator that the models do not explain the data well despite an apparently good model fit. In Experiment 3, this is the case anyways, since neither exclusive choice behavior nor behavior that would result from a stable steady-state around *c* = *c_b_* or *c* = 0 is in line with the observed response biases (see the discrepancy between the data and the black reference lines, and cp. Section 3.2).

## E Additional results for Experiment 4 and 5

Model fits and simulation behavior for example subject 9 in Experiment 4 and 5 was shown in Figures 9 and 12 respectively; for all other subjects they can be seen in Figure E.1 (Experiment 4) and Figures E.2–E.4 (Experiment 5, one figure per model).

The DT law fits to the data from Experiment 5 can be seen in Figure E.5. Note that for subject 7, in conditions 1:4 hi_2_ and 1:4 lo_2_ the data points corresponding to stimulus S1 are missing, because the subject never emitted R2 in response to this stimulus, so the log-response-ratio cannot be computed. There is no systematic difference between points from the higher-density (dark gray) and lower-density (light gray) condition of the same RR, which is in line with the conclusion from the other results of this experiment that reward density does not affect criterion setting.

**Figure C.8:**
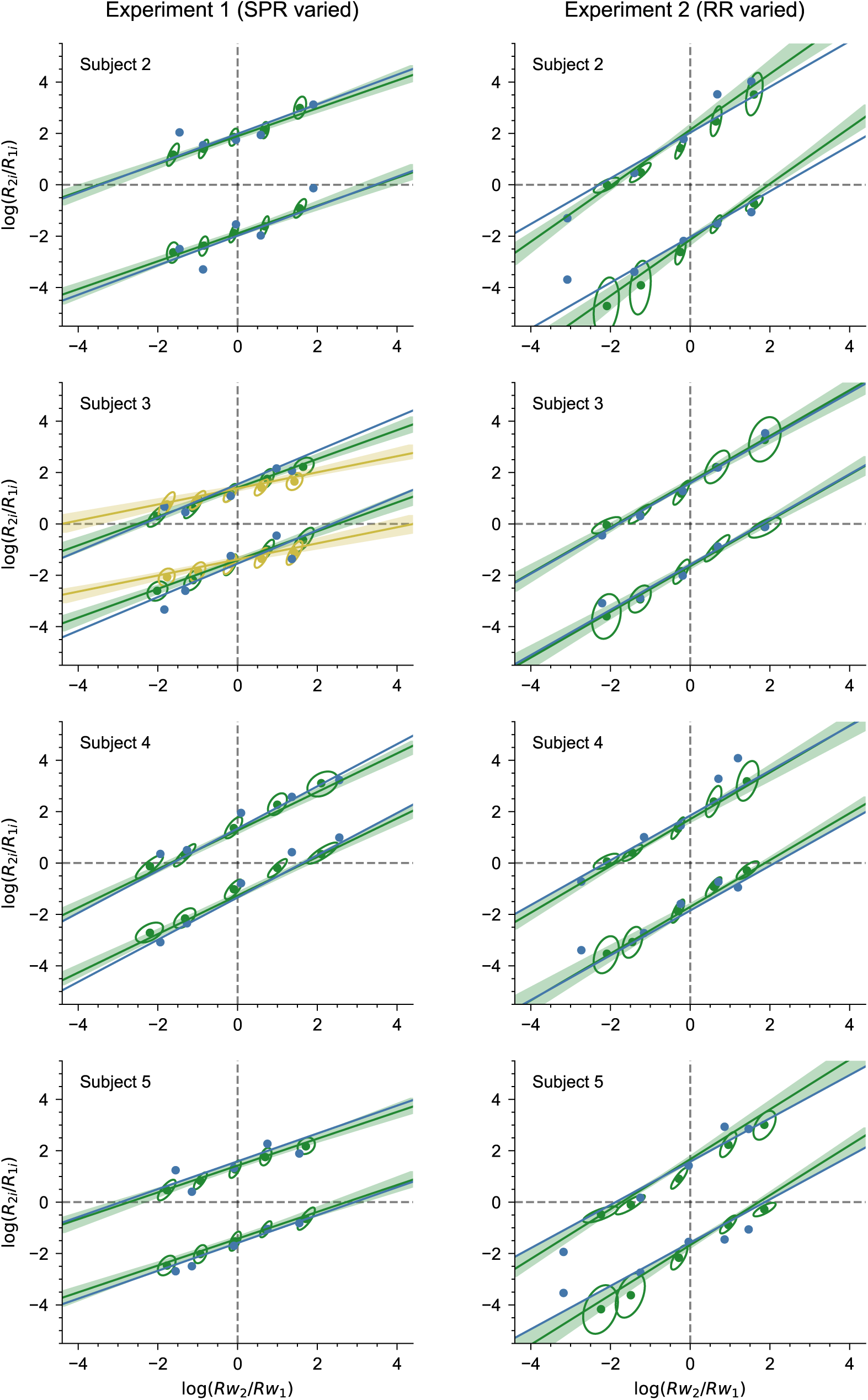
DT law fits for the modified KDB model in Experiment 1 and 2. The DT law was fitted to the data (blue) as well as to 100 simulations from the model with the best-fitting parameters (green). Like Figure C.7, left column, for the remaining subjects.

**Figure C.9:**
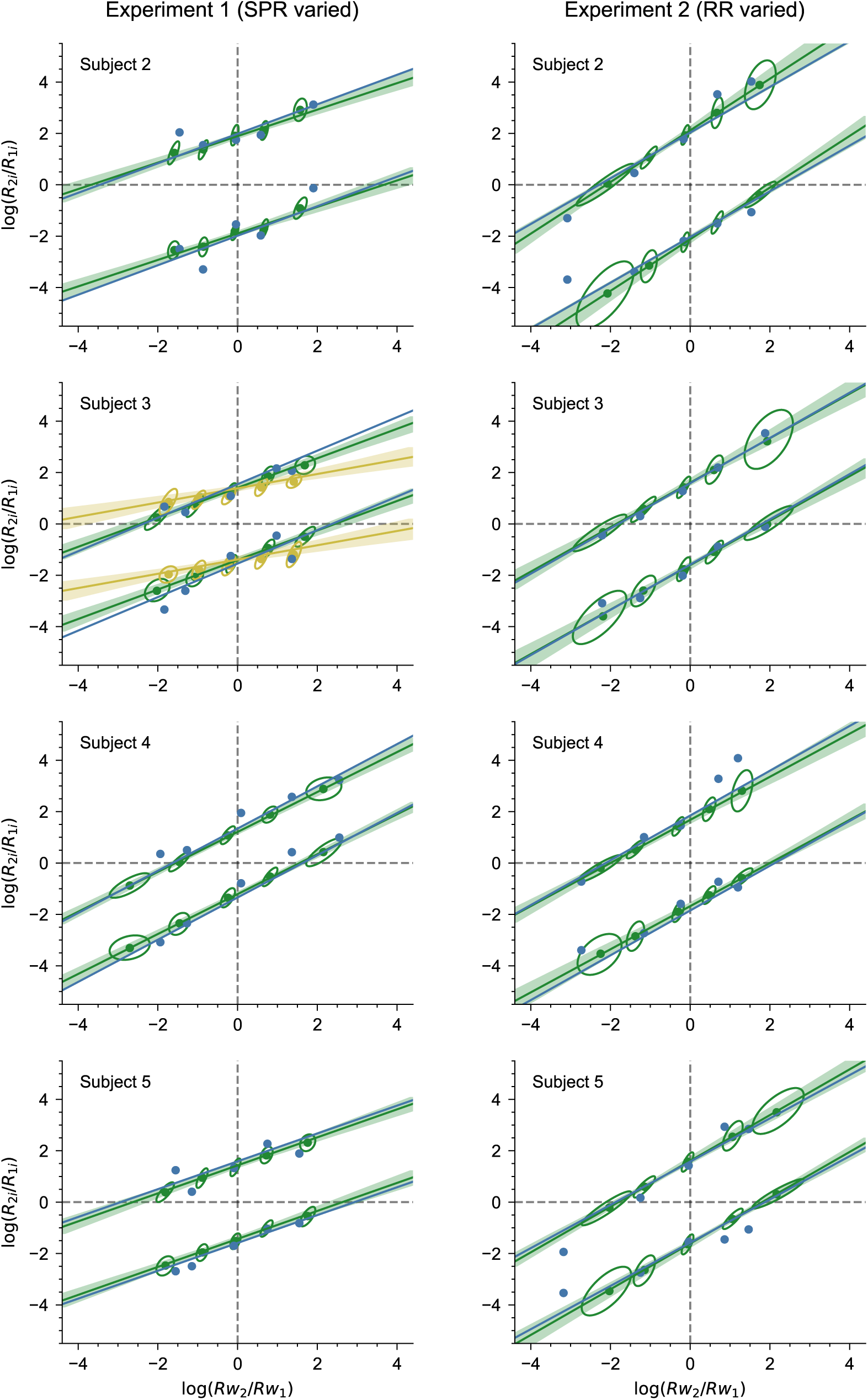
DT law fits for the DT model in Experiment 1 and 2. The DT law was fitted to the data (blue) as well as to 100 simulations from the model with the best-fitting parameters (green). Like Figure C.7, middle column, for the remaining subjects.

**Figure C.10:**
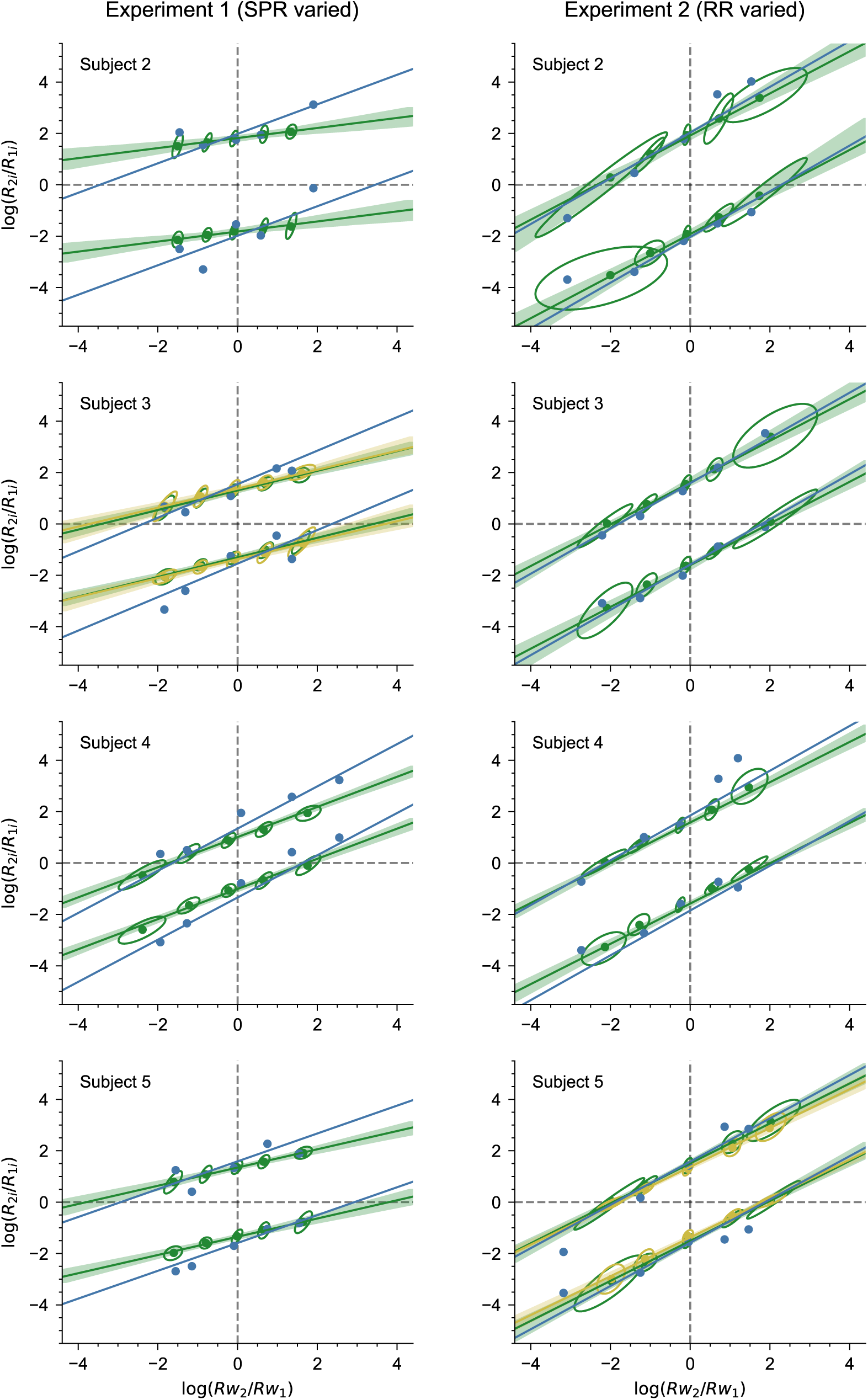
DT law fits for the RL model in Experiment 1 and 2. The DT law was fitted to the data (blue) as well as to 100 simulations from the model with the best-fitting parameters (green). Like Figure C.7, right column, for the remaining subjects.

**Figure D.1:**
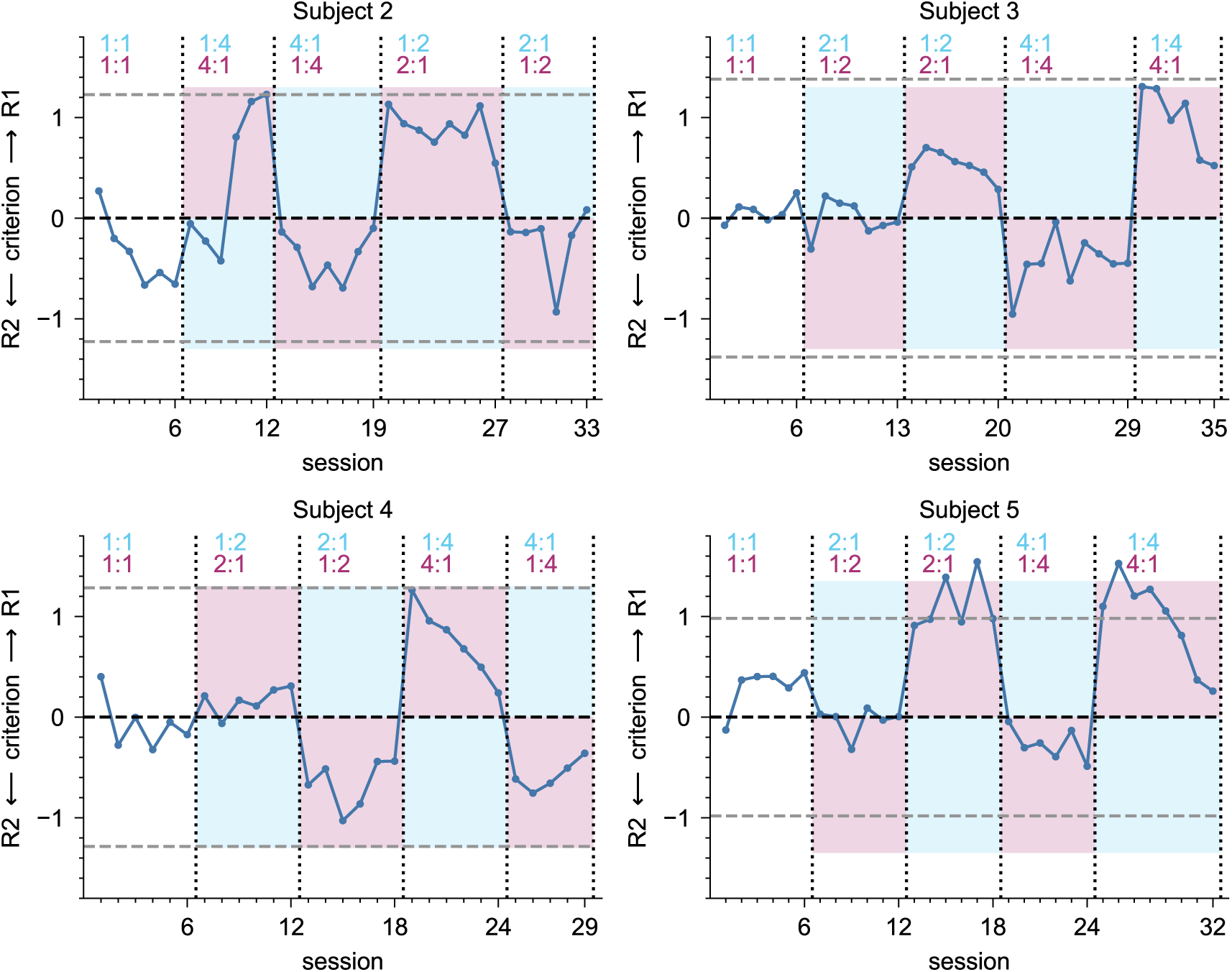
Criterion development in Experiment 3. Each data point is the criterion in one session as fitted by the one-criterion-per-session model. For reference, zero (black) and the fitted stimulus means ±*d^′^/*2 (gray) are marked as dashed lines. A positive / negative criterion indicates a bias to emit R1 / R2 respectively. Blue / purple shading indicates the areas where the criterion would indicate that SPR / RR have a stronger effect respectively. Like Figure 6B, for the remaining subjects.

**Figure D.2:**
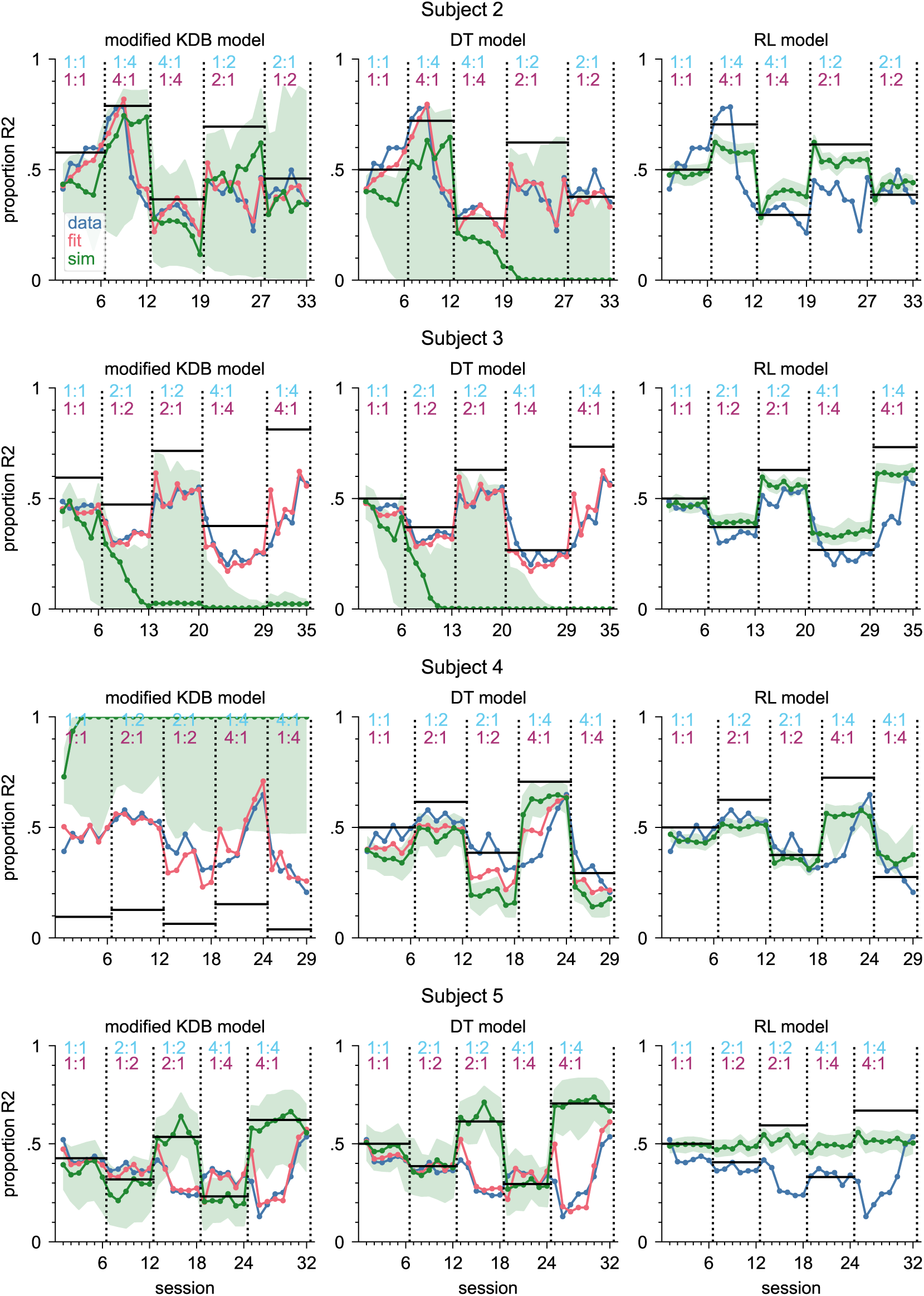
Model fit and simulations applied to the data of subjects 2-5 in Experiment 3. Each data point is the proportion of R2 in one session. Blue line: data. Red line: proportion of R2 predicted by the model fit (using the best-fitting parameters), i.e., session average of P(R2) in each trial given the actual trial history. Green line: median of 100 simulations (using the best-fitting parameters), shaded green area: central 95% of the simulations. Black lines: proportion of R2 that would result from a fixed criterion at *c_eq_* (modified KDB model) or *c* = 0 (DT and RL models) as a reference.

**Figure E.1:**
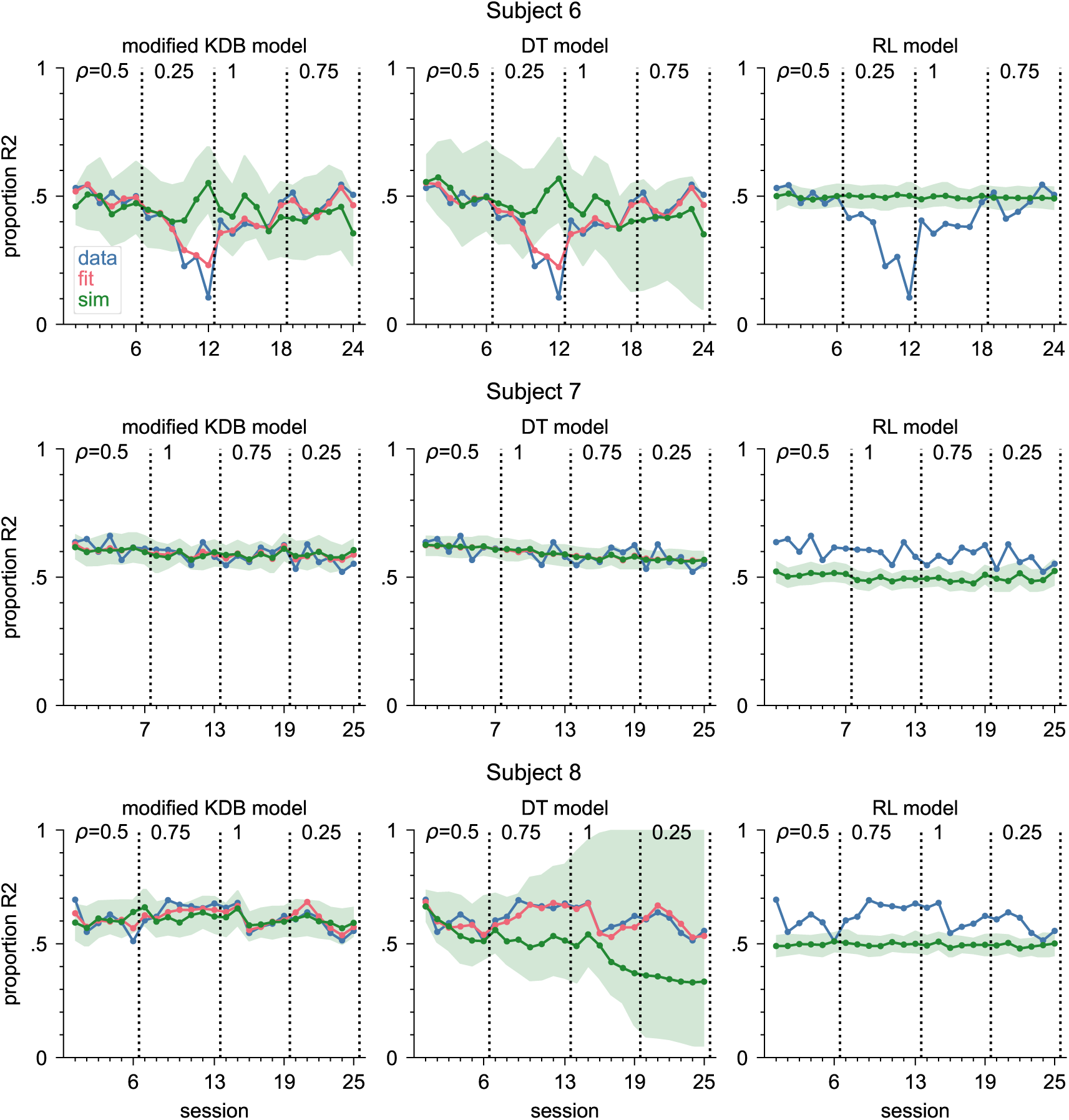
Model fit and simulations for the modified KDB model applied to the data of subjects 6-8 in Experiment 4. Each data point is the proportion of R2 in one session. Blue line: data. Red line: proportion of R2 predicted by the model fit (using the best-fitting parameters), i.e., session average of P(R2) in each trial given the actual trial history. Green line: median of 100 simulations (using the best-fitting parameters), shaded green area: central 95% of the simulations.

**Figure E.2:**
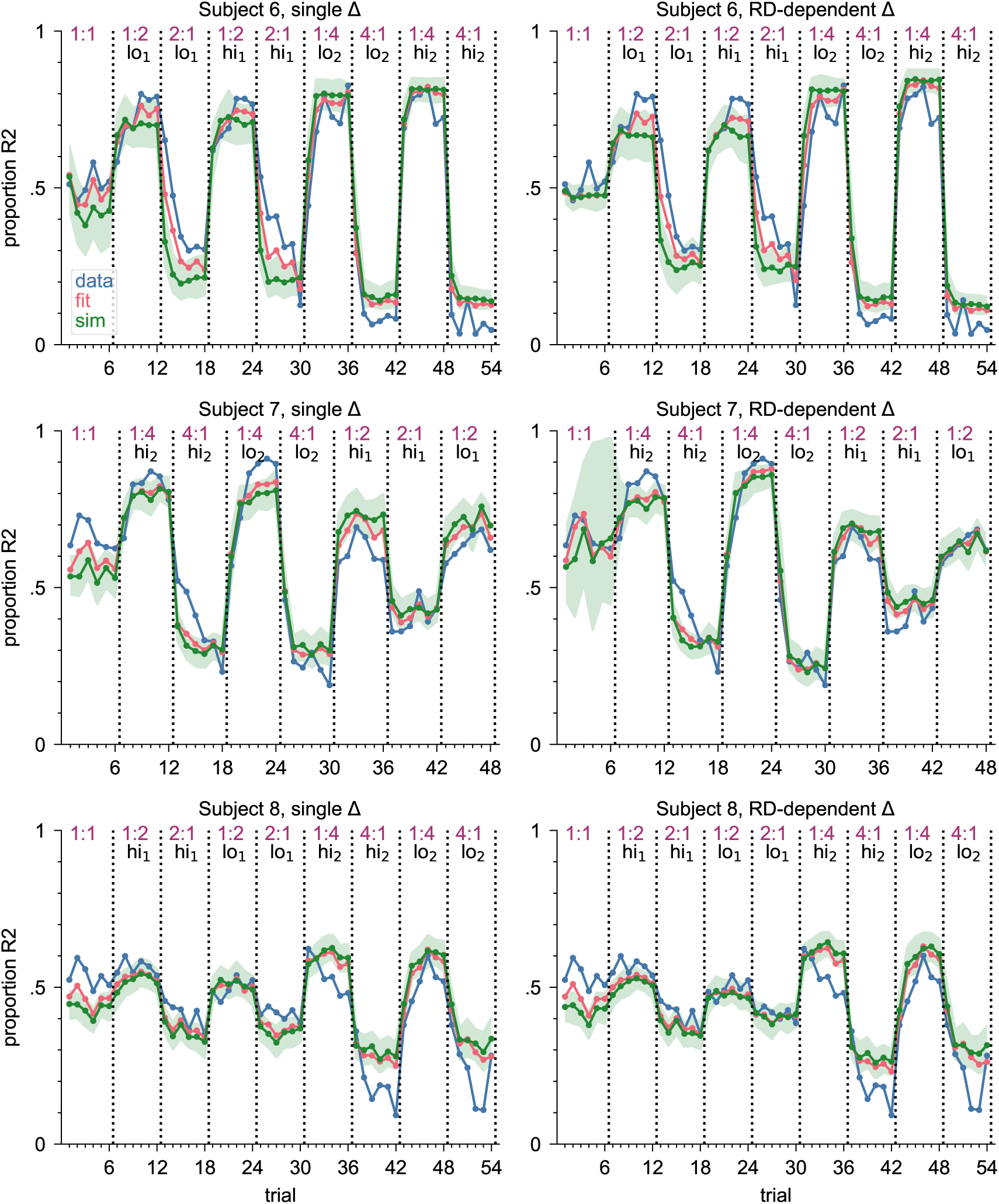
Model fit and simulations for the DT model applied to the data of subjects 6-8 in Experiment 5. Left: single-learning-rate version, right: multiple-learning-rate version. Each data point is the proportion of R2 in one session. Blue line: data. Red line: proportion of R2 predicted by the model fit (using the best-fitting parameters), i.e., session average of P(R2) in each trial given the actual trial history. Green line: median of 100 simulations (using the best-fitting parameters), shaded green area: central 95% of the simulations.

**Figure E.3:**
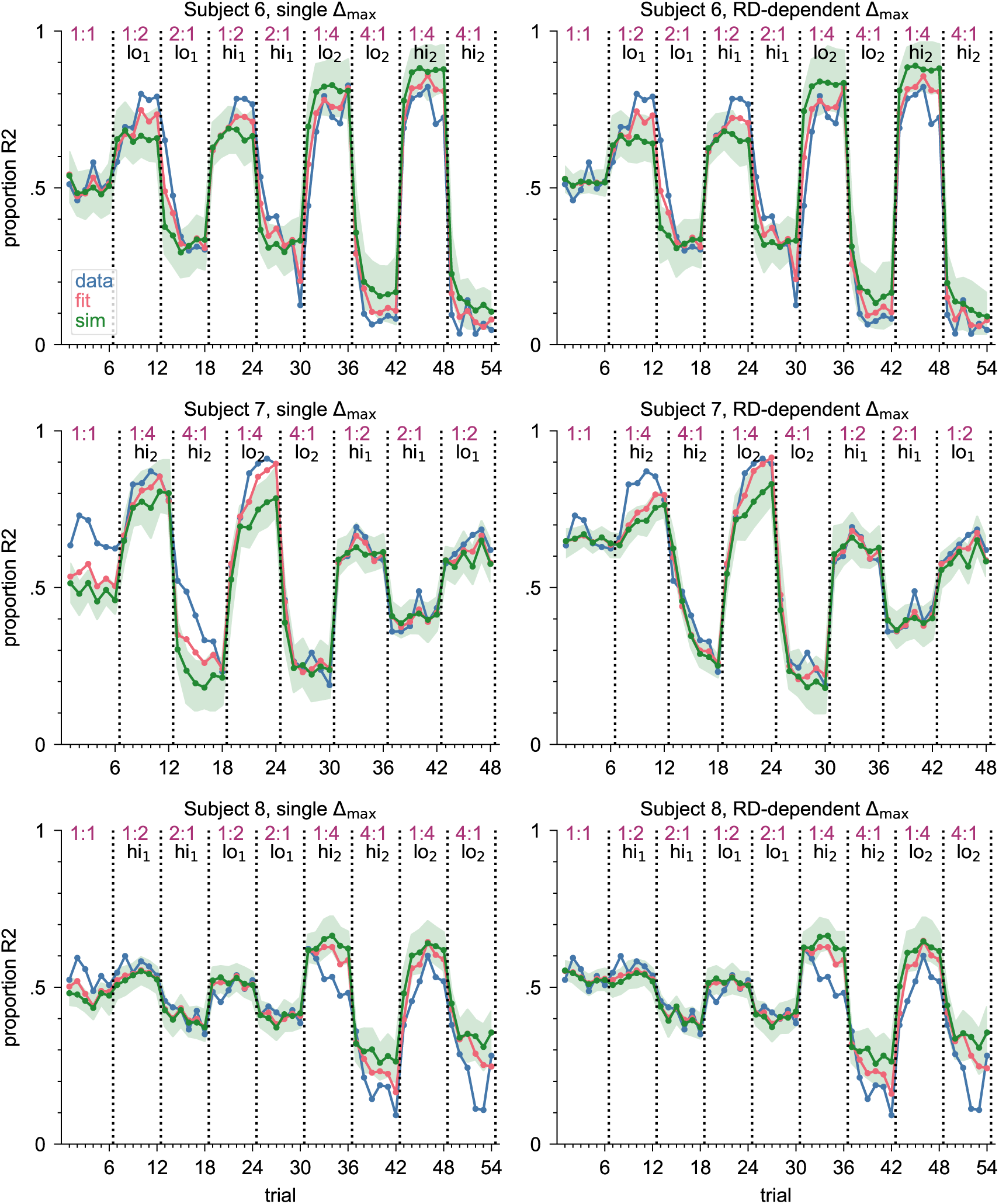
Model fit and simulations for the RL model applied to the data of subjects 6-8 in Experiment 5. Left: single-learning-rate version, right: multiple-learning-rate version. Each data point is the proportion of R2 in one session. Blue line: data. Red line: proportion of R2 predicted by the model fit (using the best-fitting parameters), i.e., session average of P(R2) in each trial given the actual trial history. Green line: median of 100 simulations (using the best-fitting parameters), shaded green area: central 95% of the simulations.

**Figure E.4:**
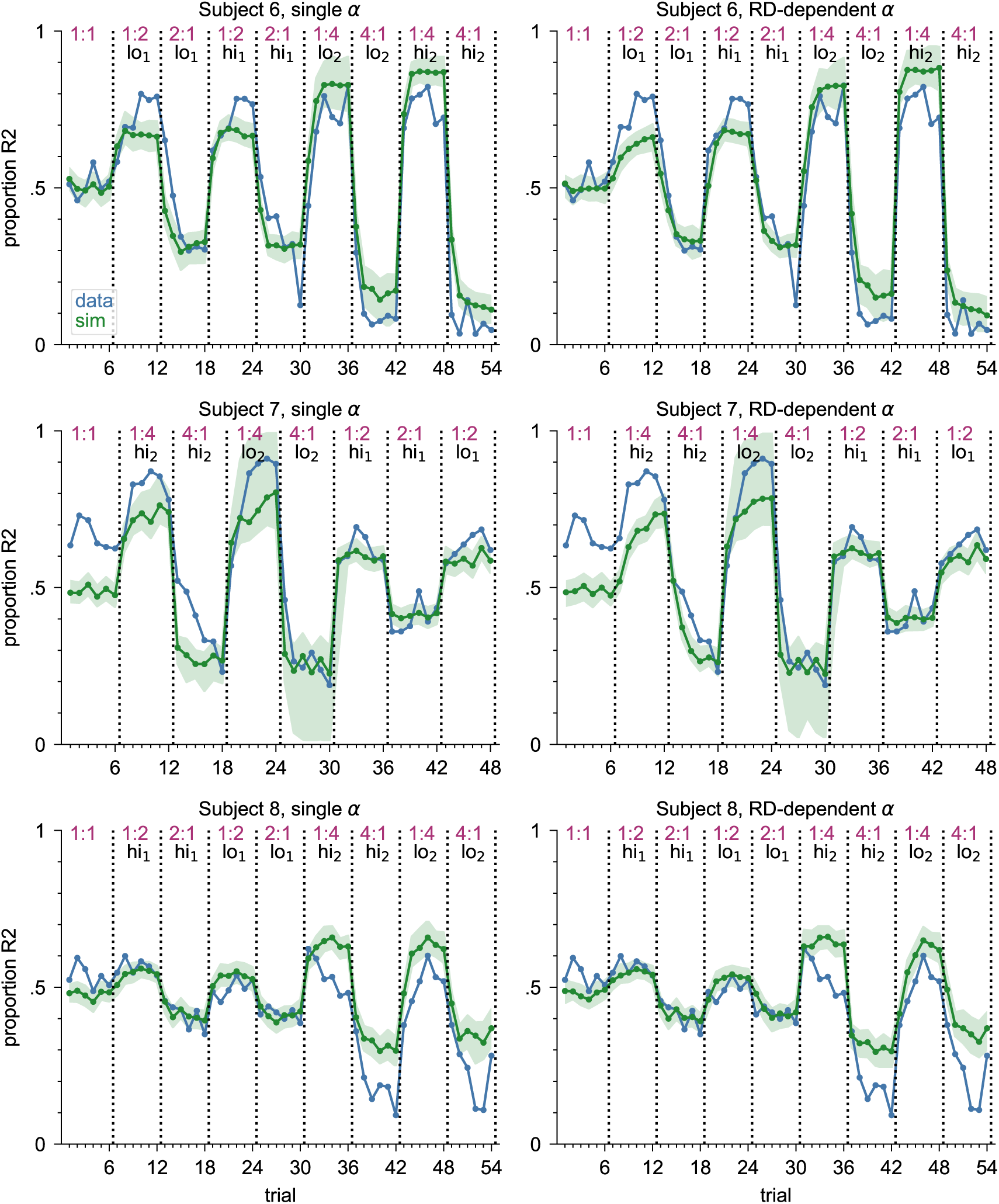
Model fit and simulations applied to the data of subjects 6-8 in Experiment 5. Left: single-learning-rate version, right: multiple-learning-rate version. Each data point is the proportion of R2 in one session. Blue line: data. Red line: proportion of R2 predicted by the model fit (using the best-fitting parameters), i.e., session average of P(R2) in each trial given the actual trial history. Green line: median of 100 simulations (using the best-fitting parameters), shaded green area: central 95% of the simulations.

**Figure E.5:**
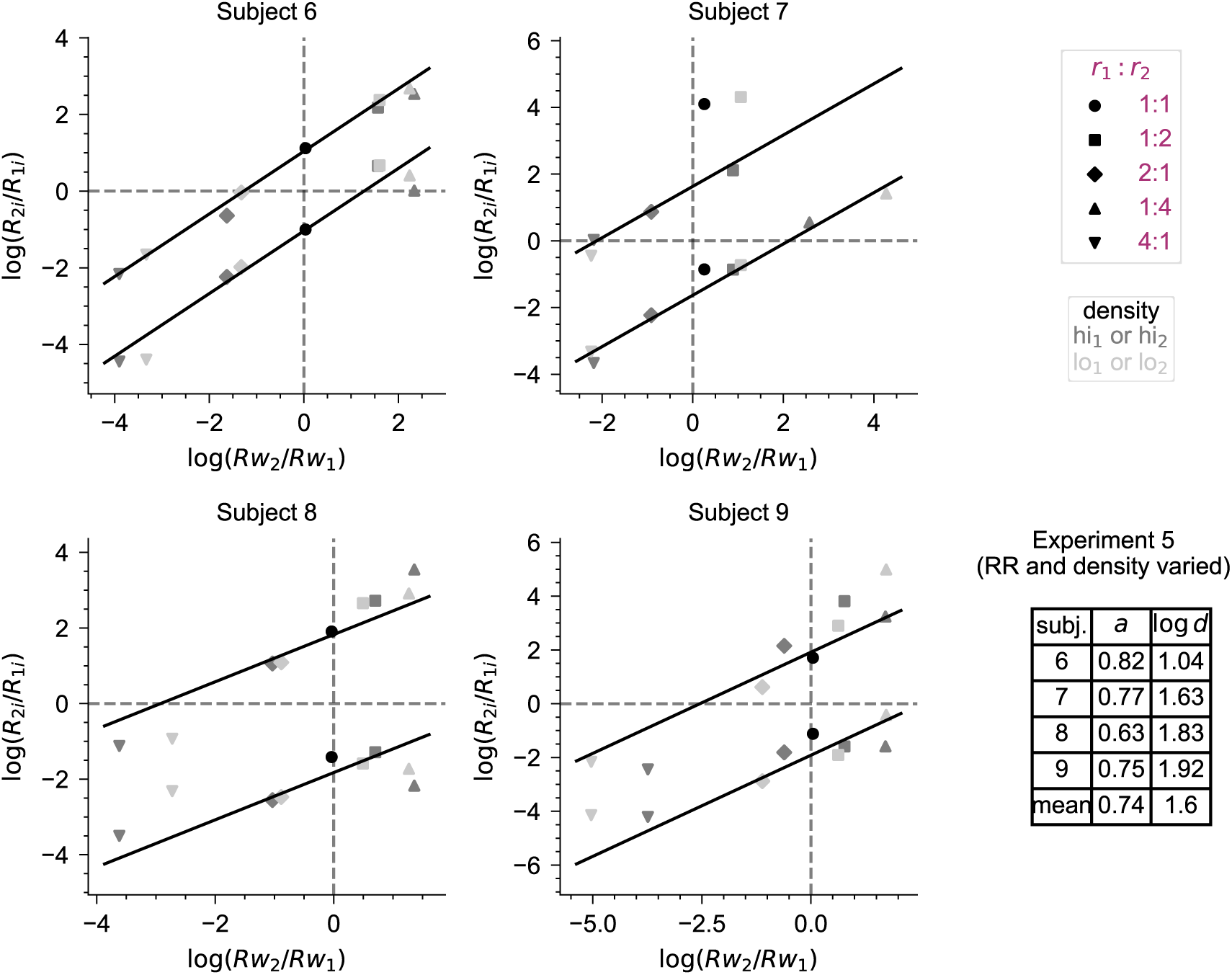
DT law fits for Experiment 5. Stimulus-wise log response ratio is plotted against reward ratio for each subject. Each data point stems from one experimental condition. Conditions that share the same RR are represented by the same shape, and data points from the hi_1_ or hi_2_ conditions and the lo_1_ or lo_2_ conditions are colored dark gray and light gray respectively, to allow an easy visual comparison between higher-density and lower-density conditions with the same RR. The lower and upper lines show the fitted linear relationship for stimulus 1 and 2 trials respectively. The fitted parameter values are shown in the bottom right corner.

## Footnotes

1 In de la Cuesta-Ferrer et al. (2025), this model is referred to as “Integrate rewards with relative differences” (IR-RD) model.

2 Note that this is exactly the IR-RD model from de la Cuesta-Ferrer et al. (2025), even though the formulas look different. In de la Cuesta-Ferrer et al. (2025), the model was parametrized to have the stimulus means *µ*_1_ and *µ*_2_ as independent free parameters (not necessarily symmetric around 0), with the baseline criterion placed at *c*(0) = 0; whereas here, the stimulus distributions are determined by a single parameter *d*^′^, with stimulus means located at −*d*^′^*/*2 and *d*^′^*/*2, and the second free parameter is the baseline criterion *c*(0) = *c_b_*. The two formulations are equivalent, but the one used here is more similar to the formulation of the DT and RL models, where the stimulus means are also centered around 0 (see Sections 2.5.2 and 2.5.3). It also has the advantage that the two model parameters here can directly be interpreted as stimulus discriminability and baseline criterion respectively.

3 *c*_0_ is the criterion position before the subject has observed anything. Therefore, it cannot be influenced by the experimental manipulations, but only reflects a response bias of the subject that is either a general preference or due to the history before the start of the experiment. Moreover, the influence of *c*_0_ on the behavior vanishes quickly over trials, as the trial-by-trial updates move the criterion position towards the equilibrium position.

4 The initial values *V*_1_(0) and *V*_2_(0) were optimistically initialized to 1 (the maximum possible reward per trial in our setup) in Experiment 1–3, while in Experiment 4 and 5 they were set to *V*_1_(0) = *V*_2_(0) = 0.5 (the most uninformed value, that assumes a reward or no reward are equally likely).

5 For subject 3 in Experiment 1, the parameters that yield the lowest NLL were very different from the other subjects’ parameters (*γ* ≪ 1 and large Δ) and did not explain the observed behavior; however, for this subject there was a second minimum in the NLL surface, with parameter values similar to the ones of the other subjects (see Figure C.1 in the appendix). We therefore used that second minimum as the “best-fitting parameters” for this subject, but still included the first one in the table in parentheses.

6 As for the modified KDB model, we also used a second minimum in the NLL as the “best-fitting parameters” for subject 3 in Experiment 1 here, because again the parameters that yield the lowest NLL in this case were very different from the other subjects’ parameters (almost zero *a* and very large Δmax) and did not explain the observed behavior, whereas those from the second minimum were more similar to the other subjects’ parameter values and explained the behavior better (see Figure C.2 in the appendix). The parameters of the first minimum are included in the table in parentheses.

7 As with the other two models, the NLL_alt_ surface had two minima for subject 3 in Experiment 1: the best-fitting one at an *α* very different from the other subjects, and another one in the same region as the other subjects’ single minimum. Additionally, this was also the case for two subjects in Experiment 2. The second minima in these cases also explained the data better (see Figures C.3 and C.6 in the appendix). Therefore, we used these as the “best-fitting parameters” for these subjects. The parameters of the first minimum for these cases are included in the table in parentheses.

8 Note that *c*_0_ always refers to the criterion at the beginning of the experiment. For each combination of 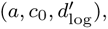 the criterion at the beginning of trial group *g_i_* is determined as the criterion at the end of the previous trial group *g_i_*_−1_ using the best 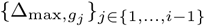 during the previous trial groups *g*_1_ to *g_i_*_−1_.

